# Chromosome-level genome assembly and methylome profile enables insights for the conservation of endangered loggerhead sea turtles

**DOI:** 10.1101/2024.08.28.610089

**Authors:** Eugenie C. Yen, James D. Gilbert, Alice Balard, Albert Taxonera, Kirsten Fairweather, Heather L. Ford, Doko-Miles J. Thorburn, Stephen J. Rossiter, José M. Martín-Durán, Christophe Eizaguirre

## Abstract

**Background:** Characterising genetic and epigenetic diversity is crucial for assessing the adaptive potential of populations and species. Slow-reproducing and already threatened species, including endangered sea turtles, are particularly at risk. Those species with temperature-dependent sex determination (TSD) have heightened climate vulnerability, with sea turtle populations facing feminisation and extinction under future climate change. High-quality genomic and epigenomic resources will therefore support conservation efforts for these flagship species with such plastic traits.

**Findings:** We generated a chromosome-level genome assembly for the loggerhead sea turtle (Caretta caretta) from the globally important Cabo Verde rookery. Using Oxford Nanopore Technology (ONT) and Illumina reads followed by homology-guided scaffolding, we achieved a contiguous (N50: 129.7 Mbp) and complete (BUSCO: 97.1%) assembly, with 98.9% of the genome scaffolded into 28 chromosomes and 29,883 annotated genes. We then extracted the ONT-derived methylome and validated it via whole genome bisulfite sequencing of ten loggerheads from the same population. Applying our novel resources, we reconstructed population size fluctuations and matched them with major climatic events and niche availability. We identified microchromosomes as key regions for monitoring genetic diversity and epigenetic flexibility. Isolating 191 TSD-linked genes, we further built the largest network of functional associations and methylation patterns for sea turtles to date.

**Conclusions:** We present a high-quality loggerhead sea turtle genome and methylome from the globally significant East Atlantic population. By leveraging ONT sequencing to create genomic and epigenomic resources simultaneously, we showcase this dual strategy for driving conservation insights into endangered sea turtles.

## BACKGROUND

With biodiversity declining at an alarming rate (Ceballos *et al*., 2015), genomic tools are being deployed to generate molecular insights for conservation management of endangered species (Theissinger *et al*., 2023). For example, characterising genetic diversity (Wold *et al*., 2021), inbreeding (Kardos *et al*., 2016), demographic history (Karamanlidis *et al*., 2021), and locally adapted genomic regions (Flanagan *et al*., 2018) can inform on the adaptive potential of populations and species (Eizaguirre and Baltazar-Soares, 2014). Conservation epigenomics has recently gained momentum, driven by increasing technological accessibility (Balard *et al*., 2024). This is a promising approach as it employs an additional layer of molecular information with a tight link to the environment (Feil and Fraga, 2012). For instance, quantifying epigenetic variation can aid in assessing a population’s capacity for adaptive plastic responses, or provide biomarkers that reflect individual health and environmental exposure (Rey *et al*., 2020; Lamka *et al*., 2022; Balard *et al*., 2024). In particular, DNA methylation — the addition of a methyl group to cytosine residues to regulate gene expression — is the best-described epigenetic modification in non-model species to date (Laine *et al*., 2023).

Sea turtles comprise seven charismatic species, with six assigned as Vulnerable to Critically Endangered and one as Data Deficient by the IUCN Red List (IUCN, 2024). Beyond threats like bycatch, poaching, and coastal development (Wallace *et al*., 2011), sea turtles are climate-vulnerable because of their temperature-dependent sex determination (TSD) system, where higher incubation temperatures induce female development (Yntema and Mrosovsky, 1982). As multiple theoretical studies have predicted complete feminisation and subsequent population collapse by 2100 in line with future climate scenarios (Hawkes *et al*., 2009; Laloë *et al*., 2014), it is essential to assess whether sea turtles can sustain viable sex ratios via adaptive responses (Mitchell and Janzen, 2010; Lockley and Eizaguirre, 2021). Although their capacity for genetic evolution is constrained by long generation times and small effective population sizes (Komoroske *et al*., 2017), plastic responses via epigenetic mechanisms could offer alternative pathways, especially as they already play a role in TSD regulation (Venegas *et al*., 2016; Ge *et al*., 2018; Piferrer, 2021). A lack of high-quality reference genomes previously hindered such molecular insights in sea turtles. However, this situation is changing through the efforts of international sequencing consortia. Chromosome-level genome assemblies have now been released for the green (*Chelonia mydas)* and leatherback (*Dermochelys coriacea*) sea turtles by the Vertebrate Genomes Project (VGP) (Bentley *et al*., 2023), and a loggerhead sea turtle (*Caretta caretta*) from the Adriatic Sea by the Canada BioGenome Project (CBP) (Chang *et al*., 2023). Yet, given the well-known influence of reference bias on downstream population-based analyses, continuing to build genome resources remains crucial (Thorburn *et al*., 2023).

Adding to the expansion of reference genomes for sea turtles, we present a chromosome-level assembly for a loggerhead sea turtle from the Cabo Verde (East Atlantic) nesting aggregation (**Figure 1A**, Population IUCN Red List Status: Endangered), which is the largest worldwide for this species (Taxonera et al. 2022). The population is composed of genetically distinct nesting groups maintained by strong female philopatry across the archipelago (Baltazar-Soares *et al*., 2020). A reference assembly for this globally important rookery will improve genomic studies by eliminating reference bias, as well as provide a high-quality loggerhead genome from the East Atlantic nesting group for comparative studies. We simultaneously present the first methylome profile derived from Oxford Nanopore Technology (ONT) reads for a sea turtle species, validated with whole genome bisulfite sequencing (WGBS) of ten loggerheads from the same population. We next applied these novel resources to describe synteny, demographic history and genomic properties. Lastly, we described the chromosomal locations of over 190 TSD-linked genes, then created a map of their methylation status and predicted functional associations, which will be a useful resource for future epigenetic studies of these endangered TSD species.

**Figure 1.**
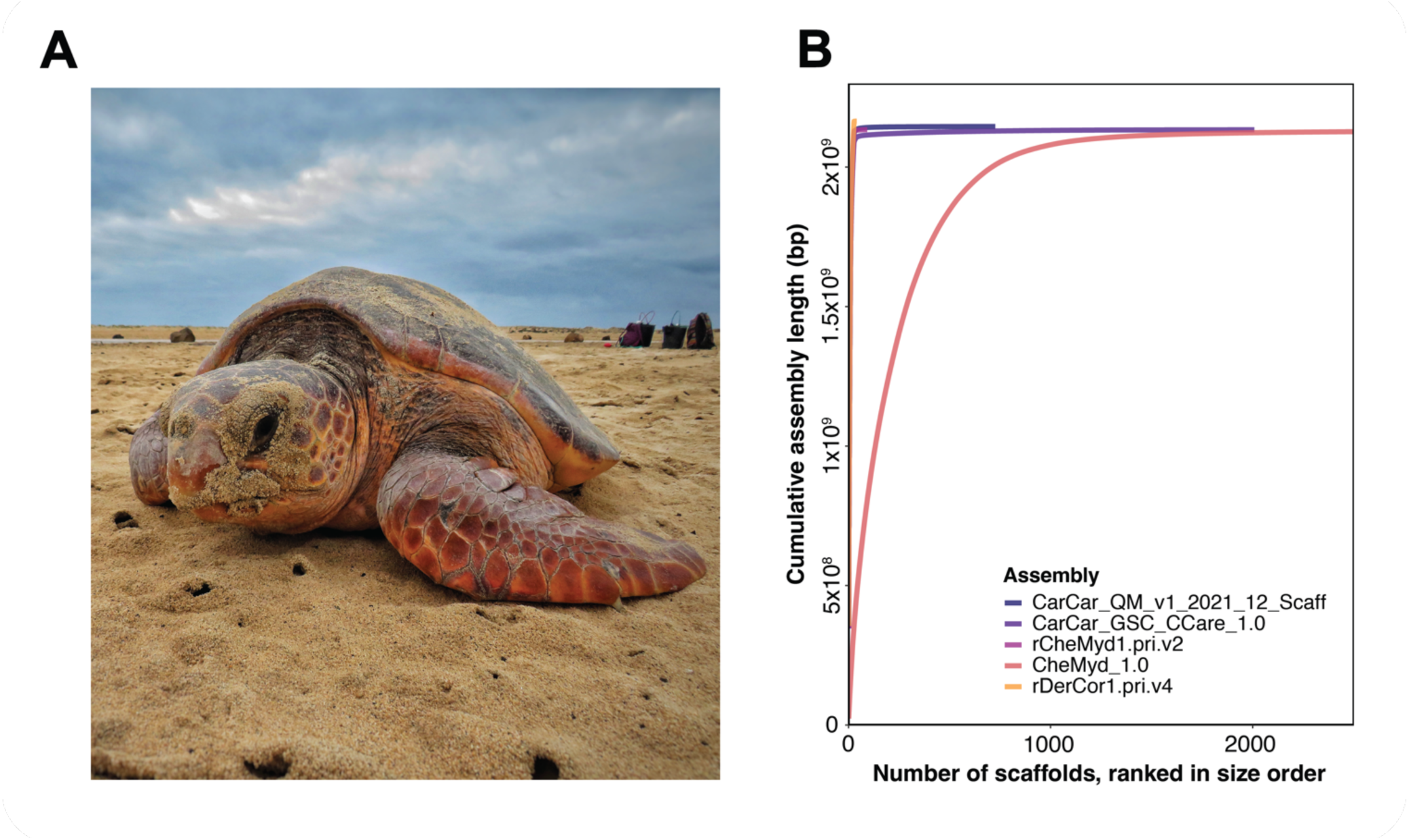
A chromosome-scale genome assembly for endangered loggerhead sea turtles from the East Atlantic nesting group. (A) A loggerhead turtle nesting in Cabo Verde (Sal Island), our reference population. Photo credit: Project Biodiversity. **(B)** Contiguity of publicly available sea turtle genomes. ‘CarCar_QM_v1_2021_12_Scaff’ (dark blue) is our chromosome-scale loggerhead (*Caretta caretta*) assembly. ‘CarCar_GSC_CCare_1.0’ (dark purple) is the chromosome-scale loggerhead assembly from the Adriatic Sea, released by the Canada BioGenome Project (Chang *et al.,* 2023) ‘rCheMyd1.pri.v2’ (mauve) and ‘rDerCor1.pri.v4’ (yellow) are chromosome-scale green (*Chelonia mydas*) and leatherback (*Dermochelys coriacea*) assemblies respectively, released by the Vertebrate Genomes Project (Bentley *et al.,* 2023). ‘CheMyd_1.0’ (pink) is the first, draft green assembly (Wang *et al.,* 2013). Our assembly is of comparable contiguity to the best genomes currently available for sea turtles.

## METHODS

### Reference sample collection

On the 31^st^ of August 2020, we sampled blood from a wild female loggerhead (ID: SLK063, Permit: 013/DNA/2020) that nested on Sal Island of the Cabo Verde Archipelago. Here, the nesting season extends between late June to October. The sampling site (16.62123 °N, - 22.92972 °E) was on Algodoeiro Beach, which consists of 800 m of sandy coastline. Blood was collected from the dorsal cervical sinus with a 40 mm, 21-gauge needle and 5 ml syringe following oviposition (Owens and Ruiz, 1980). A Passive Integrated Transponder tag was added to the front right flipper for identification (Stiebens *et al*., 2013). The sample was stored in a lithium heparin tube then centrifuged for 1 min at 3000 rpm to separate plasma and blood cells. Samples were stored at -18°C during the field season, then at -80°C following transport to Queen Mary University of London (London, UK).

### DNA extraction, sequencing and quality control

Genomic DNA was extracted from blood cells using the QIAGEN Genomic-Tips 100G Kit (Qiagen, Germany). For ONT sequencing, libraries were constructed using an SQK-LSK109 Ligation Sequencing Kit and sequencing was conducted on the PromethION 24 platform with a FLO-PRO002 flow cell (Oxford Nanopore Technologies, UK). Base calling was performed via Guppy v.4.0.11 in high-accuracy mode (Wick, Judd and Holt, 2019). This generated 11,643,721 reads (50,075,750,398 bp, ∼23.3X coverage) with an N50 of 8,230 bp (base pairs). All downstream bioinformatic steps were conducted on the Apocrita High Performance Computing Cluster (King *et al*. 2019). Adapters were trimmed with PoreChop v.0.2.4 (Wick, 2018) and reads were filtered for a Phred score >Q8 and length >500 bp with NanoFilt v.2.6.0 (De Coster *et al*., 2018). This passed 8,288,359 reads (40,371,088,741 bp, 80.6%), with an N50 of 8518 bp.

We also generated Illumina sequencing data for polishing. Libraries were constructed by fragmenting DNA via sonification, end polishing, A-tailing and adapter ligation, polymerase chain reaction (PCR) amplification with P5 and indexed P7 oligos, and purification with the AMPure XP system. Sequencing was performed with 150 bp paired end reads on the NovaSeq 6000 platform (Illumina, USA). This gave 1,050,248,476 reads (157,537,271,400 bp, ∼73.3X coverage). To aid downstream parameter selection, haploid genome size, heterozygosity and repeat content were estimated via GenomeScope (**Figure S1)** (Vurture *et al*., 2017). Reads were trimmed for adapters and filtered for a Phred score >Q20 with TrimGalore v.0.6.5 (Krueger, 2019), passing 1,050,248,476 reads (156,382,436,354 bp, 99.3%).

### *De novo* assembly

ONT reads were assembled using Flye v.2.8.3 (Kolmogorov *et al*., 2019) in ‘--asm-coverage 40’ mode. A polished consensus sequence was produced using Medaka v.1.3.3 with the ‘r941_prom_high_g4011’ model (https://github.com/nanoporetech/medaka). Using our high-coverage Illumina reads from the same sample, two rounds of error polishing were performed with Pilon v.1.24 (Walker *et al*., 2014). The contamination level was assessed via BlobTools v.1.1.1 (Laetsch and Blaxter, 2017) with Diamond BLASTx v.2.0.11 (Buchfink, Xie and Huson, 2015), comparing against the 2021_03 release of the UniProt reference proteomes database (The UniProt Consortium, 2021). The assembly was haploidised using Purge_Dups v.1.2.5 (Guan *et al*., 2020) to give a contig-level assembly “CarCar_QM_v1_2021_12”. In addition, we assembled and annotated the mitochondrial genome from our Illumina data with MitoZ v.3.4 (**Text S1** for extended methods) (Meng *et al*., 2019).

### Homology-guided scaffolding

We used the chromosome-scale loggerhead assembly produced by the CBP (Chang *et al*., 2023) for homology-guided scaffolding. Unplaced contigs were removed with SAMtools v.1.9 (Li *et al*., 2009). The remaining 28 chromosomal scaffolds served as a reference for homology-based scaffolding into chromosomes using RagTag v.2.1.0 (Alonge *et al*., 2021). We supply two versions of our assembly: (1) ‘CarCar_QM_v1_2021_12_Scaff’ with 28 chromosomes and unplaced contigs/scaffolds, and (2) ‘CarCar_QM_v1_2021_12_Scaff_With_Chr0’ where unplaced contigs/scaffolds were concatenated into a single scaffold ‘SLK063_ragtag_chr0’ with 100 bp of Ns as gap padding. Note this artificial chromosome was created as an option to aid computational analyses by reducing assembly fragmentation, and does not represent true positional information.

### Assembly quality assessment

Contiguity was evaluated with QUAST v.5.0.2 (Gurevich *et al*., 2013). Completeness was assessed with Benchmarking Universal Single-Copy Ortholog (BUSCO) scores, using BUSCO v.5.1.2 (Simão *et al*., 2015) in genome mode against the ‘sauropsida_odb10’ database (n=7480 BUSCOs). K-mer-based assessments were performed by comparing k-mers from our Illumina reads against the assembly with parameter K=21. A k-mer spectrum was produced using KAT v.2.4.1 (Mapleson *et al*., 2017) and an assembly consensus quality value (QV) was calculated with Merqury v.1.3 (Rhie *et al*., 2020). Contiguity and completeness comparisons were also conducted against all assemblies available for sea turtles: (1) ‘CarCar_GSC_CCare_1.0’ produced by the CBP for the Adriatic loggerhead turtle (Chang *et al*., 2023), (2) ‘rDerCor1.pri.v4’ and (3) ‘rCheMyd1.pri.v2’ produced by the VGP for the leatherback and green turtle respectively (Bentley *et al*., 2023), and (4) ‘CheMyd_1.0’ which was the first draft assembly for the green turtle (Wang *et al*., 2013).

### Genome annotation

A repeat library was built using RepeatModeler v.2.0.4 in ‘LTRStruct’ mode to discover lateral terminal repeats (Flynn *et al*., 2020). To exclude potential *bona fide* gene families, the repeat library was compared against the proteome of the VGP green sea turtle (Bentley *et al*., 2023) via Diamond BLASTp v.2.0.11 (Buchfink, Xie and Huson, 2015). Transposable elements (TEs) were classified using TEclass (Abrusán *et al*., 2009), then the curated repeat library was used for annotation via RepeatMasker v.4.1.4 (Smit *et al.,* 2022).

For gene annotation, paired-end RNA-Seq reads (n=746,132,735, **Table S1**) from 24 loggerheads across three life stages (hatchling, juvenile and adult), four tissue types (blood, gonad, brain and heart) and both sexes were mined from the Sequence Read Archive (Leinonen, Sugawara and Shumway, 2011; Banerjee *et al*., 2021; Chow *et al*., 2021; Hernández-Fernández *et al*., 2021). Reads were trimmed with Trimmomatic v.0.36 (Bolger, Lohse and Usadel, 2014), mapped with STAR v.2.7.10a (Dobin *et al*., 2013), and sorted with SAMtools v.1.9 (Li *et al*., 2009). Alignments were then supplied to BRAKER1 for gene prediction (Hoff *et al*., 2016). Note that species-specific training was attempted but resulted in a poorer annotation. Chicken (*Gallus gallus domesticus*) pre-trained parameters were hence chosen, as they were the most related species available. ProtHint v.2.6.0 (Brůna *et al*., 2021) was used to prepare protein hints from OrthoDB’s odb10 database of Vertebrata protein sequences (Kriventseva et al. 2019), as well as predicted proteomes of the VGP green and leatherback sea turtle (Bentley *et al*., 2023). These were supplied to BRAKER2 for gene prediction. TSEBRA v.1.0.3 (Gabriel *et al*., 2021) was then implemented in ‘pref_braker1’ mode to combine BRAKER1 and BRAKER2 outputs into a set of best predictions.

Concurrently, gene prediction followed the Mikado pipeline v.2.2.4 (Venturini *et al*., 2018) with whole transcriptome and transcript-based hints. Transcriptomes included a blood transcriptome produced from eight loggerheads across three life stages (Hernández-Fernández *et al*., 2021) and a transcriptome we assembled *de novo* using Trinity v.2.14 (Grabherr *et al*., 2011) with RNA-Seq data (**Table S1**) for gonad, brain and heart tissue of three hatchlings (Chow *et al*., 2021). Transcriptomes were mapped to our assembly via GMAP v.2021-12-17 (Wu and Watanabe, 2005), with a 99.56% and 99.98% alignment rate, respectively. From our STAR alignments, curated intron junctions were produced with Portcullis v.1.2.3 (Mapleson *et al*., 2018) and open reading frames were calculated using TransDecoder v.5.5.0 (Haas, 2018). All evidence types were subsequently supplied for gene prediction by Mikado.

The BRAKER and Mikado gene sets were merged using the PASA pipeline v.2.5.2 (Haas *et al*., 2008) with three rounds of comparison. Finally, the gene set was filtered and standardised with AGAT v.0.9.1 (Dainat, 2022), and in-frame stop codons were removed with gffread v.0.12.7 (Pertea *et al*., 2015). To assess completeness, BUSCO v.5.1.2 (Simão et al. 2015) was run on the longest isoforms in protein mode against the ‘sauropsida_odb10’ database (n=7480 BUSCOs). Homology-based functional information was assigned via Diamond BLASTp v.2.0.11 (Buchfink, Xie and Huson, 2015) against the SwissProt database v.2022_03_02 (The UniProt Consortium, 2021). Gene Ontology (GO) terms were added using InterProScan 5 v.5.60-92.0 with HMMER databases: Gene3D-4.3.0, PANTHER-17.0, Pfam-35.0, PIRSR-2021, SFLD-4, SUPERFAMILY-1.75, TIGRFAM-15.0 (Jones *et al*., 2014). All functional annotations were attached via MAKER v. 2.31.9 (Cantarel *et al*., 2008).

### ONT methylation call and validation with WGBS

From our ONT reads, we called 5-methylcytosine (5mC) and 5-hydroxymethylcytosine (5hmC) modifications in the CpG (5’-C-phosphate-G-3) context via Guppy v.6.5.7 (Wick, Judd and Holt, 2019) in configuration ‘dna_r9.4.1_450bps_modbases_5hmc_5mc_cg_hac_prom’. We focused on CpGs as this is the main methylation context in vertebrates (Klughammer *et al*., 2023). As methylation also occurs symmetrically at CpG sites in vertebrates (Klughammer *et al*., 2023), calls were de-stranded per CpG then converted to bedMethyl files using Modkit v.0.1.9 mpileup (https://github.com/nanoporetech/modkit). This retained 26,449,075 CpGs.

To validate that our ONT-derived methylation call was comparable to a gold-standard method, we generated WGBS-derived methylomes of ten nesting loggerheads from the same population and locality as the reference individual (**Table S3**), using the same sampling protocol. Genomic DNA was extracted with a QIAGEN Blood and Tissue Kit (Qiagen, Germany). DNBseq libraries were constructed and sequenced with 100 bp paired-end reads on an MGI DNBSEQ platform (BGI, Hong Kong), generating 132,192,522 ± 40,360 (SD) reads per sample (**Table S4**). Full methods for methylation calling are available in **Text S2.** Briefly, alignment and methylation calling were performed via Bismark v.0.22.1 (Krueger and Andrews, 2011), with 79.1 ± 3.37 (SD) % mapping efficiency. CpGs were de-stranded using the ‘merge_CpG.py’ script (Cristofari, 2022), resulting in a coverage of 9.2 ± 0.33 (SD) X (**Table S4**). Using the R package methylKit v.1.24.0 (Akalin *et al*., 2012), CpGs were filtered for coverage between 5X and the 99.9^th^ percentile (Wreczycka *et al*., 2017), then retained if they were covered in >75% of individuals, leaving 24,299,151 CpGs. For each CpG, a methylation percentage was calculated per individual with methylKit’s ‘percMethylation’ function. The mean across all individuals provided an average WGBS methylome representative of the population.

To compare ONT and WGBS methylation calls, 5mC and 5hmC modifications were merged for the ONT data because WGBS cannot distinguish between them. As methylation patterns are known to differ between feature types (Jones, 2012), we calculated mean methylation across four categories (promoter, exon, intron and intergenic regions; extended methods in **Text S3**) with the R packages genomation v.1.30.0 (Akalin *et al*., 2015) and GenomicRanges v.1.50.2 (Lawrence *et al*., 2013). A chi-squared test was used to evaluate if highly methylated (>70%) CpGs (Sagonas *et al*., 2020) were distributed differently across feature types between ONT and WGBS data. Secondly, a linear model was used to test if mean methylation per gene was correlated between ONT and WGBS data, interacting by feature type. The gene-associated feature types included were promoters, exons, introns and intergenic regions <10 kbp (kilo base pairs) from the nearest transcription start site (TSS) (Heckwolf *et al*., 2020). All statistical analyses were conducted in R v.4.2.2 (R Core Team 2021) and all plots were produced with the R package ggplot2 v.3.4.2 (Wickham, 2016).

### Genome-wide synteny between sea turtle species

To investigate genome-level synteny between sea turtle species, we mapped our loggerhead assembly against the VGP green and leatherback assemblies (Bentley *et al*., 2023). Genomes were aligned using minimap2 v.2.18-r1015 with parameter ‘-f 0.02’ (Li, 2018) and dot plots were produced in D-GENIES v.1.5.0 (Cabanettes and Klopp, 2018).

### Demographic history

We performed Pairwise Sequentially Markovian Coalescent (PSMC) analysis (Li and Durbin, 2011) to reconstruct the effective population size (Ne) of the Cabo Verdean loggerhead population (East Atlantic), using our Illumina reads for the reference individual SLK063 (Morin *et al*., 2021). To extend insights across the Atlantic Ocean, we repeated this analysis with Illumina reads for a loggerhead from a Brazilian (Bahia, West Atlantic) population (BioSample: SAMN20502673, SRA Run: SRR15328383) (Vilaça *et al*., 2021). Reads were aligned via BWA-MEM v.0.7.17 (Li, 2013), with a mapping rate of 99.8% for SLK063 and 98.7% for SAMN20502673. Alignments were sorted with SAMtools v.1.9 (Li *et al*., 2009), and duplicates were tagged with Picard MarkDuplicates v.2.26.9 (Broad Institute, 2022). Variant calling and consensus building were conducted in BCFtools v.1.19 with a base and mapping filter of >Q30 (Danecek *et al*., 2021). Sites between a third and twice the mean coverage (SLK063: ∼51.7X, SAMN20502673: ∼22.5X) were retained (Bentley *et al*., 2023). PSMC v.0.6.5 was run on the eleven macrochromosomes (1.73 Gbp, 80.8% of total assembly) with parameters ‘-N25 -t15 -r5 -b -p “4+25*2+4+6’ (Bentley *et al*., 2023) and 100 bootstraps over ∼17 million years ago (Mya). PSMC was also run on microchromosomes to verify that patterns were similar (**Figure S2)**. Outputs were scaled with a mutation rate of 1.2^-8^ (Bentley *et al*., 2023) and a generation time of 45 years. This was calculated by adding the age of maturity to half the reproductive longevity (Vilaça *et al*., 2021), where the age of maturity was estimated as ∼30 years in Cabo Verde using the length-at-age relationship (Piovano *et al*., 2011). Global mean surface temperature anomaly was plotted relative to pre-industrial times with climate data inferred from marine sediments (Hansen *et al*., 2013; Clark *et al*., 2024).

### Genome properties

We used our Illumina data for the reference individual to estimate genome-wide heterozygosity (Robinson *et al*., 2019). Reads were aligned via BWA-MEM v.0.7.17 with a mapping rate of 99.6% (Li, 2013), sorted with SAMtools v.1.9 (Li *et al*., 2009), and PCR duplicates were tagged with Picard MarkDuplicates v.2.26.9 (Broad Institute, 2022). Variants were called including monomorphic sites with GATK v.4.2.6.1 HaplotypeCaller in ‘-ERC BP_RESOLUTION’ mode, followed by genotyping via GenotypeGVCFs with expected heterozygosity set to 0.00179, as estimated by GenomeScope (**Figure S1**) (McKenna *et al*., 2010). Unused alternate alleles were removed, and sites between a third and twice the mean coverage (∼56.1X) were retained (Bentley *et al*., 2023). Other filters applied were quality by depth >2.0, root mean square mapping quality >50.0, mapping quality rank sum test >−12.5, read position rank sum test >−8.0, Fisher strand bias <60.0 and strand odds ratio <3.0. With the remaining 2,086,358,400 sites, heterozygosity was computed in non-overlapping 100 kbp windows with the ‘popgenWindows.py’ script in ‘-indHet’ mode (Martin, 2016).

Next, we summarised a selection of genetic and methylation properties per chromosome in our reference assembly, to explore differences between the 11 macrochromosomes and 17 microchromosomes of the loggerhead genome (Machado *et al*., 2020). Genetic properties included were mean heterozygosity, gene density (number of genes by chromosome length) and CpG density (number of CpG sites by chromosome length). Methylation properties included were mean methylation and proportion of highly methylated (>70%) CpGs. Wilcoxon rank-sum tests were used to investigate mean differences between chromosome types per property. To examine relationships between properties, separate linear models were implemented to test whether different genome properties were correlated, with an interaction by chromosome type to test if relationships differed between macro- and microchromosomes.

### TSD-linked genes: identification and synteny between sea turtle species

To identify TSD-linked genes in our loggerhead assembly (**Text S4** for extended methods), we used a list of 223 genes compiled by Bentley et al. (2023). These genes have documented links to TSD, primarily from studies of freshwater turtles and alligators (Bentley *et al*., 2023). Following manual curation of this list, we identified the longest isoform orthologues of VGP green turtle sequences (Bentley *et al*., 2023) in our loggerhead genome via BLASTn v.2.11.0 (Altschul *et al*., 1990). These were manually verified by integrating BLAST output and gene name information. For loggerhead genes that matched sequences on two chromosomes, both locations were retained if they were syntenic with other species, under the assumption of a conserved duplication across sea turtles. If one sequence was syntenic and the other not, the non-syntenic sequence was removed under a conservative assumption of assembly error. This gave 201 unique TSD-linked genes for downstream analyses. We next assessed the chromosomal locations of TSD-linked genes in our loggerhead assembly against the VGP green and leatherback assemblies. After filtering for genes present in all three species, 199 genes across 205 loci were retained for comparison and plotted with the R package Circlize v.0.4.16 (Gu *et al*., 2014).

### TSD-linked genes: comparison of methylation patterns

We examined whether methylation differs between TSD-linked and non-TSD-linked genes in the loggerhead blood methylome. For the non-TSD-linked gene set, we used single-copy orthologues representing evolutionarily conserved genes in sea turtles. These were identified with OrthoFinder v.2.5.4 (Emms and Kelly, 2015) in nucleotide mode between our loggerhead, VGP green, and VGP leatherback genomes (Bentley *et al*., 2023), excluding TSD-linked genes. Overall, 200 unique TSD-linked genes and 15,041 single-copy orthologues were covered in our loggerhead methylome and included in analyses. We tested whether methylation differs between gene categories. To satisfy statistical assumptions, 1000 random subsamples of 200 orthogroups were generated for comparison against the 200 TSD-linked genes. For each subsample, a linear model was used to test if mean methylation was associated with gene category, with an interaction by feature type. A quasipoisson generalised linear model was used to test whether the highly methylated CpG count was associated with gene category in an interaction by feature type, with an offset of total CpG count.

### TSD-linked genes: a functional association map

We built a functional association network for TSD-linked genes with the STRING v.12.0 database (Szklarczyk *et al*., 2020). Protein sequences from the VGP green turtle (Bentley *et al*., 2023) were available on STRING and used as query sequences. Proteins were retained if they matched the name of the target gene or had a sequence match >80%. This left 191 proteins, which were searched with the following options: full STRING network, 0.4 confidence and 5% false discovery rate stringency. The Markov Clustering algorithm was used to identify clusters within the network with inflation parameter 2.2. Promoter methylation status in our loggerhead genome was annotated onto the protein network per TSD-linked gene, with three categories based on the bimodal distribution observed: high (>70%), low (<30%) and intermediate (30-70%) methylation. A chi-squared test was used to investigate if proportions of promoter methylation categories differed between the five largest clusters.

## RESULTS AND DISCUSSION

### Genomic resources generated

#### Genome assembly

By combining long ONT and short Illumina read sequencing, we produced a contig-level assembly (**Table 1**) with a total size of 2.146 Gbp, 1,799 contigs and an N50 of 5.51 Mbp (Mega base pairs). A blob-plot confirmed minimal contamination, with 99.2% of contigs mapping to Chordata and the remainder yielding no taxonomic hits (**Figure S3).** We also assembled and annotated the mitochondrial genome from our Illumina reads, consisting of a circular 16,574 bp contig with 37 genes (**Text S1, Figure S4**). Following homology-guided scaffolding against the same species (Chang *et al*., 2023), 98.9% (2.123 Gbp) of the assembly was placed into 28 chromosomal scaffolds, with 698 unplaced contigs (22.7 Mbp; 1.06% of assembly). This elevated our assembly to chromosome-level contiguity with an N50 of 129.73 Mbp, comparable to the best assemblies available for sea turtle species (**Figure 1B**, **Table 1**). With a BUSCO completeness score of 97.1% (Single copy: 96.2%, Duplicated: 0.9%, Fragmented: 0.4%, Missing: 2.5%) our assembly is the most complete among sea turtles after the VGP green genome (Bentley *et al*., 2023), and the most complete loggerhead genome to date (**Table 1; Table S4** for BUSCO summaries). Quality is further supported by a QV score of Q41.1 (>99.99% assembly accuracy) and a k-mer spectrum indicating successful haploidisation (**Figure S5)**. Overall, these comparisons demonstrate that our loggerhead assembly is a high-quality contribution to sea turtle genome resources.

**Table 1.** Comparison of assembly quality metrics across sea turtle species. BUSCO scores (full summary in Table S4) were calculated against the Sauropsida gene set (n=7480) with BUSCO v.5.1.2 (Simão *et al*., 2015). ‘CarCar_QM_v1_2021_12_Scaff’ is our chromosome-scale loggerhead (*Caretta caretta*) assembly. ‘CarCar_GSC_CCare_1.0’ is the chromosome-scale loggerhead assembly released by the Canada BioGenome Project (Chang *et al*., 2023) ‘rCheMyd1.pri.v2’ and ‘rDerCor1.pri. are chromosome-scale green (*Chelonia mydas*) and leatherback (*Dermochelys coriacea*) assemblies respectively, released by the Vertebrate Genomes Project (Bentley *et al*., 2023). ‘CheMyd_1.0’ is the first, draft green assembly (Wang *et al*., 2013).

| Assembly | Size (Gb) | Total scaffold / contig count | Longest scaffold / contig (Mbp) | N50 (Mbp) | N50 count | GC content (%) | Complete BUSCOs (%) |
| --- | --- | --- | --- | --- | --- | --- | --- |
| **CarCar_QM_v1_2021_12**<br>(Loggerhead turtle, contig) | 2.146 | 1799 | 24.43 | 5.51 | 118 | 43.94 | 97.1% |
| **CarCar_QM_v1_2021_12_Scaff**<br>(Loggerhead turtle, scaffolded) | 2.146 | 726 | 352.82 | 129.73 | 5 | 43.94 | 97.1% |
| **CarCar_GSC_CCare_1.0**<br>(Loggerhead turtle, scaffolded) | 2.134 | 2008 | 345.74 | 130.96 | 5 | 44.03 | 96.1% |
| **rDerCor1.pri.v4**<br>(Leatherback turtle, scaffolded) | 2.165 | 41 | 354.45 | 137.57 | 5 | 43.35 | 96.3% |
| **rCheMyd1.pri.v2**<br>(Green turtle, scaffolded) | 2.134 | 93 | 348.27 | 134.43 | 5 | 44.01 | 97.2% |
| **CheMyd_1.0**<br>(Green turtle, contig) | 2.132 | 140,023 | 22.92 | 4.07 | 149 | 43.43 | 95.9% |

#### Genome annotation

We identified and masked 924.6 Mbp (43.1% of assembly) of repetitive elements (**Table S5**). A total of 29,883 genes were annotated, of which 23,690 genes (82.0%) were functionally annotated via homology and 16,954 genes (58.7%) were assigned GO terms (**Table S6**). Our annotation had a BUSCO completeness score of 94.7% (Single copy: 93.9%, Duplicated: 0.8%, Fragmented: 1.1%, Missing: 4.2%). This is less complete than existing sea turtle annotations (**Table S7**), likely due to the use of sub-optimal parameters for BRAKER training (Hoff *et al*., 2019; Brůna *et al*., 2021). Future improvement could involve species-specific tuning and manual curation steps.

#### Methylation call

By calling methylation from our ONT reads, we provide an additional layer of molecular information to facilitate epigenomic insights for sea turtle conservation (Balard *et al*., 2024). In total, 22,327,230 CpGs had only 5mC modifications, 120,983 CpGs had only 5hmC modifications, and 2,986,762 CpGs had a combination of both. To verify our ONT methylation call, we compared calls with ten loggerhead methylomes re-sequenced via WGBS, the current gold-standard for base-resolution methylation analysis (Olova *et al*., 2018). The proportion of highly methylated CpGs was 74.1% for the ONT methylome versus 73.9% for the WGBS methylome. Methylation estimates were therefore consistent between sequencing methods, as well as a reported genome-wide methylation level of ∼70% across Testudines (Klughammer *et al*., 2023). Highly methylated CpGs were also similarly distributed across feature types between sequencing methods (χ^2^=0.0387, p=0.998, **Figure 2A**). For 24,884 genes covered by both methods (98.1% of ONT gene set, 98.2% of WGBS gene set), mean methylation per gene was strongly and positively correlated between ONT and WGBS (R^2^=0.92, t=1007.2, p<0.0001), with an interaction by feature type, as expected (WGBS methylation x Feature type: F_1,3_=48.9, p<0.0001, **Figure 2B**). The demonstrated comparability with WGBS supports ONT as a robust alternative approach for methylation analysis (Simpson *et al*., 2017; Xu and Seki, 2020). By measuring real-time ionic current fluctuations, ONT enables the simultaneous acquisition of genomic and methylation sequencing data from native DNA. This maximises molecular insights whilst minimising technical biases introduced by amplification and bisulfite conversion. Furthermore, ONT can distinguish between epigenetic modification types which have different regulatory implications, such as 5mC and 5hmC (Shen and Zhang, 2013).

**Figure 2.**
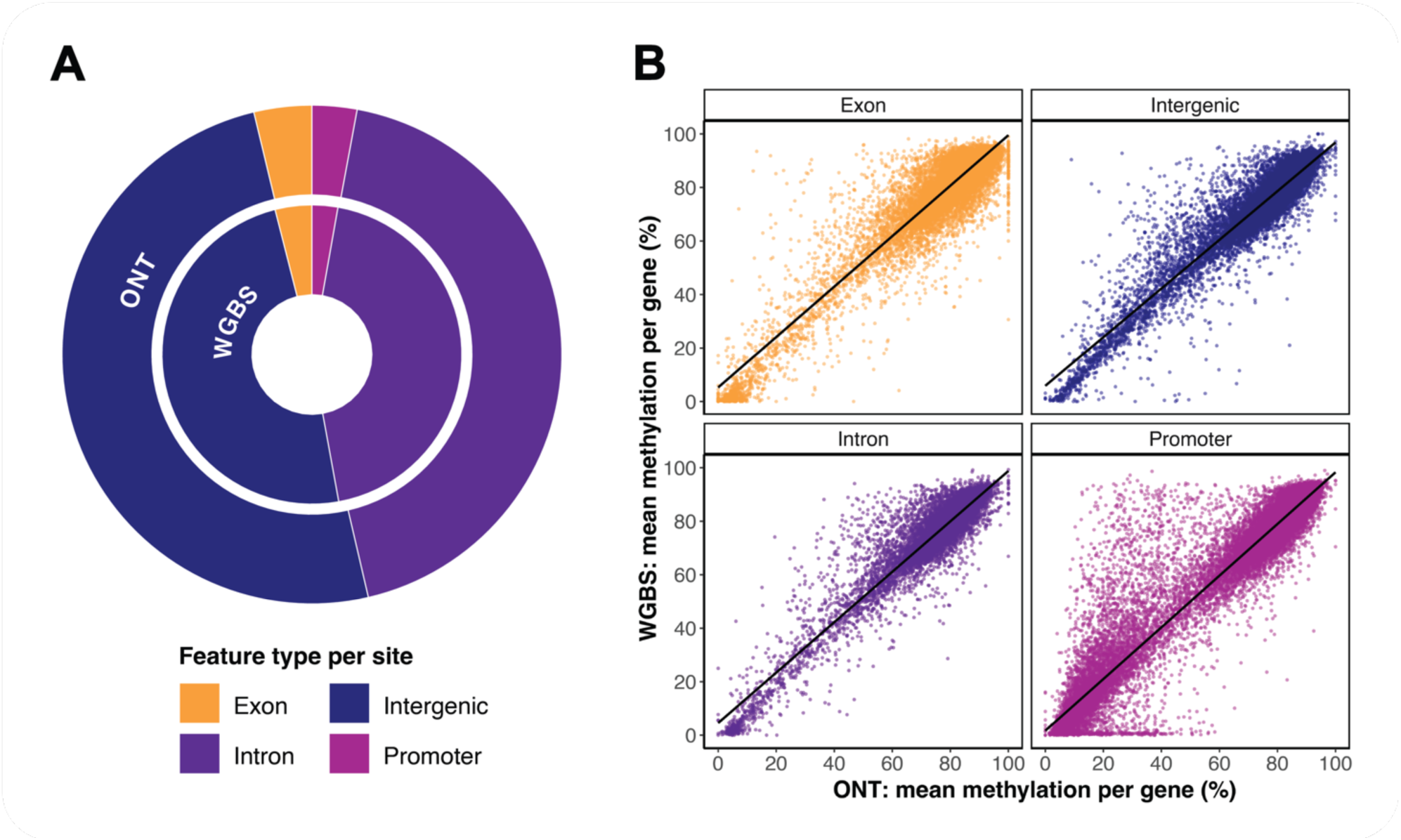
Validation of ONT-derived methylome with WGBS of ten loggerheads. (A) Genome-wide distribution of highly methylated (>70%) CpGs across feature types by site. The outer ring shows the distribution called from the ONT methylome of the reference individual (n=19,606,231 CpGs), with 3.74% on exons (yellow), 24.64% on introns (dark purple), 2.90% on promoters (mauve) and 49.89% being intergenic (dark blue). The inner ring shows the distribution called from the average of ten WGBS methylomes (n=17,950,597 CpGs), with 3.96% on exons, 24.96% on introns, 2.78% on promoters and 49.91% being intergenic. Highly methylated CpGs are similarly distributed across feature types between sequencing methods over the entire genome (χ2=0.0387, p=0.998). (B) Correlation of mean methylation (%) per gene (n=24,884) between ONT and average WGBS methylation calls, by feature type: exons, introns, promoters and gene-associated intergenic regions (<10 kbp from TSS). There is a strong, positive correlation between sequencing methods for genes per feature type (R^2^=0.92, t=1007.2, p<0.0001).

### Application of genomic resources

#### Genome-wide synteny between sea turtle species

The loggerhead genome was highly syntenic against the green (**Figure 3A)** and leatherback genomes overall (**Figure 3B**). Possible inversions were detected on chromosomes 4 and 9 of the loggerhead genome against both species, which should be checked to verify if they are true loggerhead-specific rearrangements or assembly artefacts. 90.1% of sequences exhibited >50% similarity and 2.81% lacked a match against the green, versus 3.37% with >50% similarity and 7.57% without a match against the leatherback (**Table S8**). This aligns with phylogenetic expectations, as loggerheads and greens are both members of the Cheloniidae family and diverged ∼40 Mya, compared to the deeper split ∼75 Mya from leatherbacks in the Dermochelyidae family (Vilaça *et al*., 2021). Nonetheless, our results extend the observation of high genomic stability in the sea turtle lineage (Bentley *et al*., 2023).

**Figure 3.**
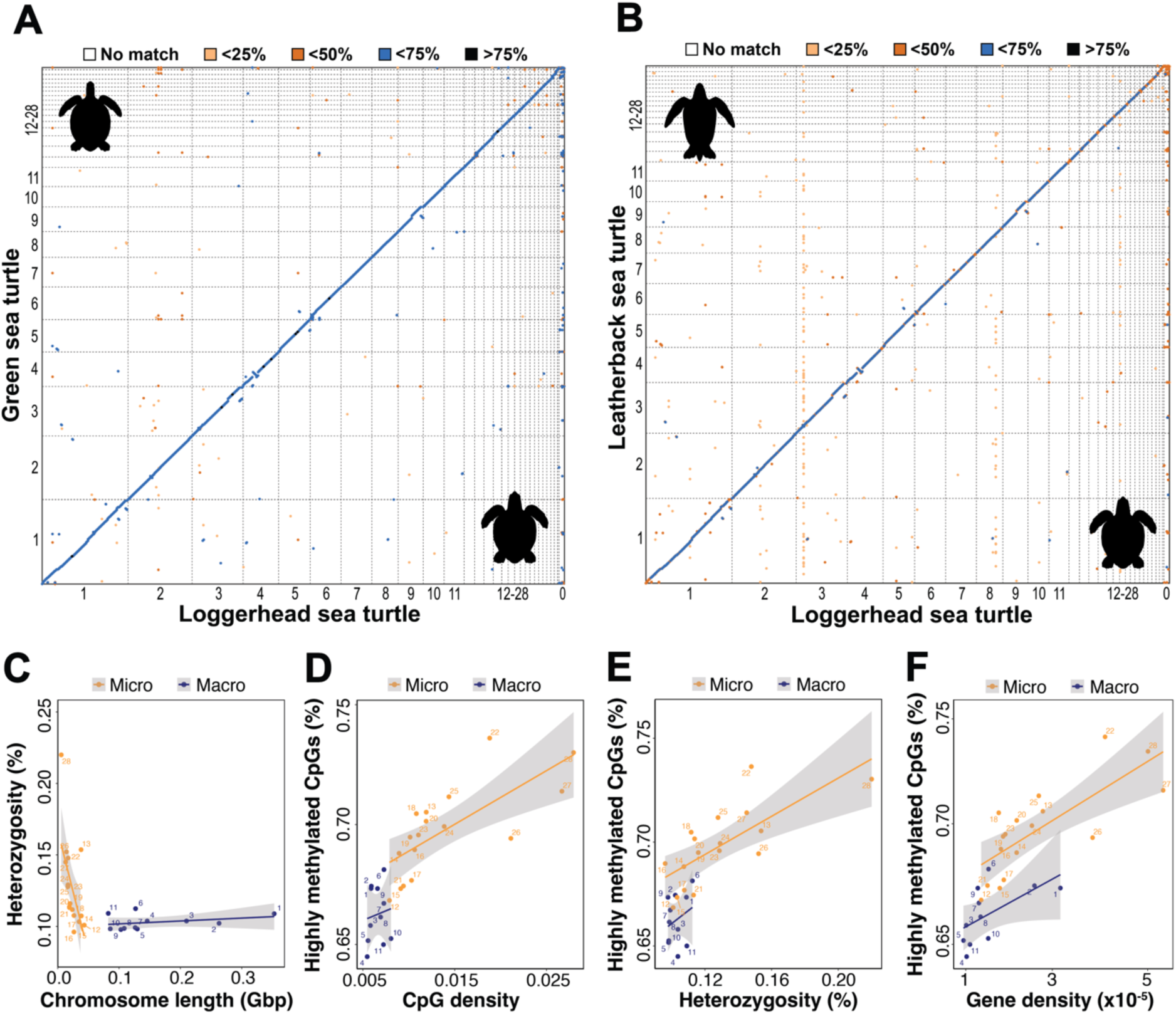
Comparison of genome-level and chromosome-level characteristics. (A-B) Alignment dotplots of our loggerhead assembly against the Vertebrate Genomes Project **(A)** green assembly and **(B)** leatherback assembly (top 100,000 alignments shown). Axes show chromosome number (1-11: macrochromosomes, 12-28: microchromosomes, 0: unplaced contigs). Colours represent alignment sequence identity (%). There is strong synteny with minimal chromosome-scale structural rearrangements between sea turtle species, supporting high genomic stability. **(C-F)** Relationships between genome properties in macro- (dark blue) versus microchromosomes (yellow) of the loggerhead genome. **(C)** Mean heterozygosity (%) exhibits an interaction by chromosome length (Gbp) and type (F_1,24_=12.3, p=0.002), with a negative correlation in microchromosomes but no correlation in macrochromosomes. **(D)** The proportion of highly methylated (>70%) CpGs (%) is higher overall for microchromosomes, but the positive correlation with CpG density (total CpGs by chromosome length; F_1,24_=56.5, p<0.0001) does not interact with chromosome type (F_1,24_=0.08, p=0.90). A similar relationship is observed between highly methylated CpGs (%) and **(E)** mean heterozygosity (%; (F_1,24_=50.30, p<0.0001, Interaction: F_1,24=_0.04, p=0.84), and **(F)** gene density (total genes by chromosome length x10^-5^; F_1,24_=65.89, p<0.0001, Interaction: F1,24=0.048, p=0.83). This suggests that relationships between methylation and other genetic properties are a genome-level property unaffected by chromosome type.

#### Demographic history

We used PSMC to reconstruct the effective population size (Ne) of loggerheads from East Atlantic (ID: SLK063, Cabo Verde) and West Atlantic (ID: SAMN20502673, Brazil, Bahia State; Vilaça et al 2021) populations. An overall decline in Ne was detected across ∼17 Mya of reconstruction (**Figure 4**). In line with contemporary estimates, the East Atlantic population had a higher Ne (∼5000) than the West Atlantic population (∼2000) near present times, although the West Atlantic population had higher Ne over the populations’ histories. Interestingly, Ne fluctuations were similar in timing and amplitude for both populations. This suggests that major climatic and oceanic processes affecting the entire Atlantic Ocean, rather than region-specific events, were the primary drivers of loggerhead demographic changes. Specifically, a population contraction occurred during the onset and intensification of Northern Hemisphere Glaciation (∼3.3-2.4 Mya), as global temperatures and atmospheric carbon dioxide dropped and ice sheets expanded (Martínez-Botí *et al*., 2015; McClymont *et al*., 2023). This period of cooling, coupled with the migration of high productivity centres from polar to equatorial regions, likely reduced habitable zones for loggerheads, as reflected in the relatively rapid decrease in Ne during this interval. During the mid-Pleistocene Transition (∼1.25-0.60 Mya), prolonged and intensified glacial intervals resulted in glacial cooling and expansion of ice sheets (Ford and Chalk, 2020). At that time, productivity centres shifted from equatorial regions toward subpolar regions and were increasingly variable on glacial-interglacial time scales (Lawrence *et al*., 2013; Lamy *et al*., 2024). These shifts may coincide with Ne expansion in loggerheads, and also matches their proposed migration history in the Atlantic Ocean (Baltazar-Soares *et al*., 2020). Altogether, our results emphasise the importance of climate and niche availability for sea turtle population dynamics, which can inform predictive models of demographic responses to current anthropogenic climate change.

**Figure 4.**
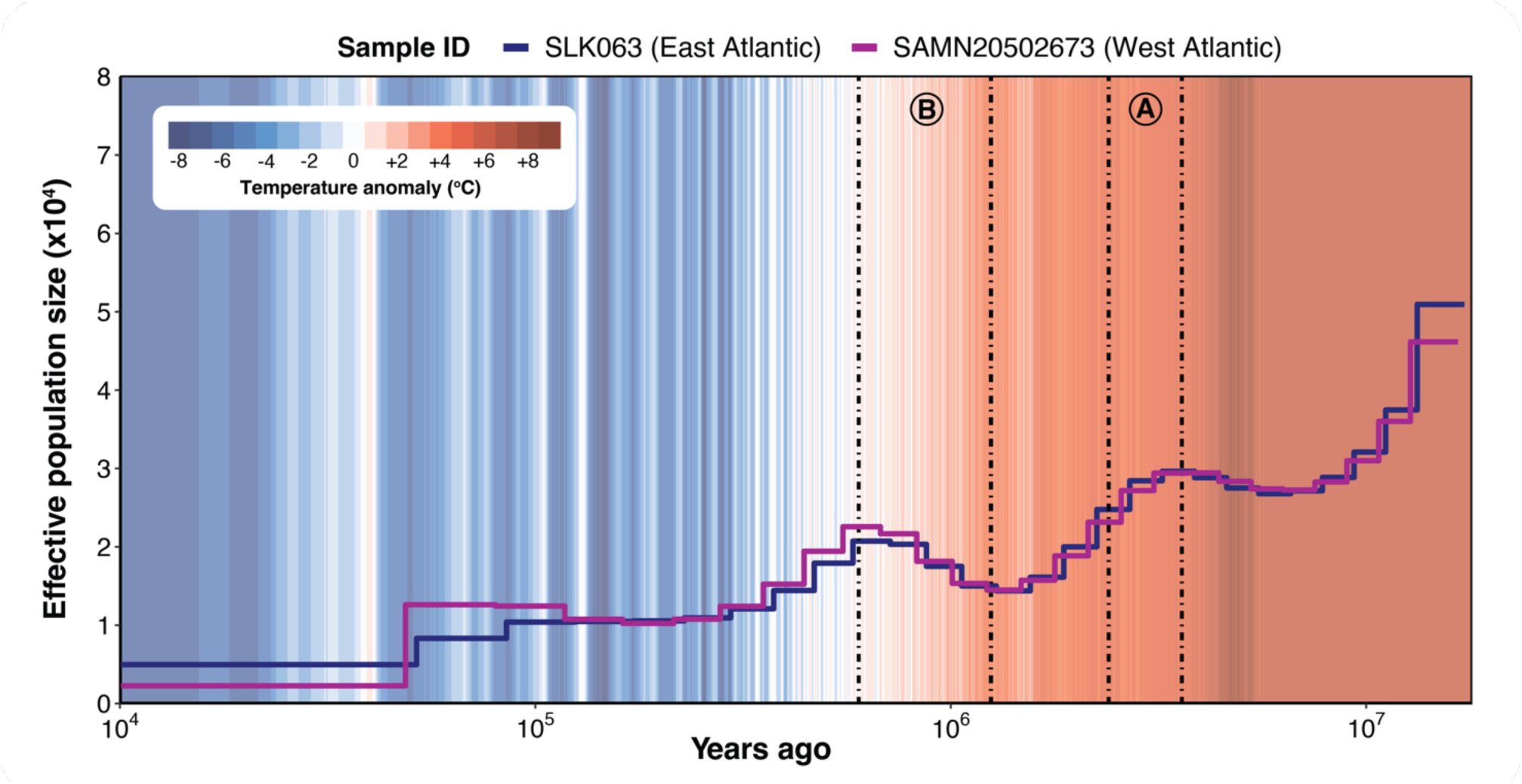
Demographic history reconstruction for loggerhead populations across the East and West Atlantic Ocean. The East Atlantic population is in Cabo Verde (SLK063, blue line) and the West Atlantic population is in Bahia State, Brazil (SAMN20502673, purple line; Vilaça, et al., 2021). Effective population size (Ne, x104) was reconstructed using PSMC over ∼17 Mya (log-scaled). Marine sediments were used to infer the global mean surface temperature anomaly (red and blue climate stripes) compared to pre-industrial times (Hansen, et al., 2013; Clark, et al., 2024). Letters designate geoclimatic events of interest: A: Northern Hemisphere Glaciation (∼3.3 to 2.4 Mya). During this period, global temperature and atmospheric carbon dioxide decreased and ice sheets expanded (Martínez-Botí, et al., 2015; McClymont, et al., 2023). High productivity centres shifted from polar to equatorial regions (K. T. Lawrence, et al., 2013), likely forcing migration and shrinkage of ecological niches. B: Mid-Pleistocene Transition (∼1.25 to 0.60 Mya). A period of glacial cooling and ice sheet growth (Ford and Chalk, 2020) when productivity centres shifted from the equator toward subpolar regions (Lawrence et al., 2013; Lamy et al., 2024), allowing population expansion.

#### Genome properties

Genome-wide heterozygosity was estimated as ∼0.12% for our loggerhead genome, in alignment with ∼0.11% reported for an Adriatic loggerhead (Chang *et al*., 2023). This places the genomic diversity of loggerheads at approximately four times the heterozygosity of leatherbacks (∼0.0029%) and half of greens (∼0.25%) (Bentley *et al*., 2023), likely reflecting their effective population sizes. Genomic diversity further varied across the 28 chromosomes, with microchromosomes being more heterozygous than macrochromosomes (W=29, p=0.002, **Figure S6A**). Heterozygosity was best predicted by an interaction between chromosome length and type (Length x Type: F_1,24_=12.3, p=0.002, **Figure 3C)**, with a steep negative correlation in microchromosomes (F_1,15_=15.4, p=0.001) but none detected for macrochromosomes (F_1,15_=4.50, p=0.063). Microchromosomes were also more gene-dense (W=28, p=0.001, **Figure S6B**), GC-rich (W=1, p<0.0001, **Figure S6C**) and CpG-dense (W=1, p<0.0001, **Figure S6D**), consistent with patterns described in other sea turtles (Bentley *et al*., 2023) and wider vertebrates (McQueen *et al*., 1996; Waters *et al*., 2021).

We next performed comparisons to address the knowledge gap of methylation differences between macro- and microchromosomes. Mean methylation was similar between chromosome types (W=54, p= 0.07, **Figure S6E**), but microchromosomes had a greater proportion of highly methylated CpGs (W=54, p<0.0001; **Figure S6F**). This likely stems from higher CpG density on microchromosomes providing more raw genetic substrate for methylation (Papin *et al*., 2021), as supported by a strong, positive correlation (F_1,24_=56.5, p<0.0001, **Figure 3D**) that was independent of chromosome type (CpG density x Type: F_1,24_=0.08, p=0.90). Similar relationships were observed for highly methylated sites against heterozygosity (F_1,24_=50.30, p<0.0001, Heterozygosity x Type: F_1,24_=0.04, p=0.84; **Figure 3E**) and gene density (F_1,24_=65.89, p<0.0001, Gene density x Type: F_1,24_=0.048, p=0.83; **Figure 3F**). Therefore, microchromosomes have greater methylation potential and realised levels than macrochromosomes, with correlations between methylation and other genome properties likely shaped by evolutionary forces that act consistently across chromosome types. These results highlight the specific properties of vertebrate microchromosomes (Waters *et al*., 2021) and add novel insights into their methylation potential for sea turtles. With higher heterozygosity, gene density and regulatory opportunities via CpG methylation, microchromosomes could serve as hotspots for both adaptive evolution and plastic responses to the environment. As such, microchromosomes offer promising large-scale regions for monitoring and preserving functional genetic and epigenetic diversity in sea turtles (Eizaguirre and Baltazar-Soares, 2014; Hoelzel, Bruford and Fleischer, 2019).

#### TSD-linked genes: synteny between sea turtle species

As TSD species, loggerheads are particularly vulnerable to climate change, with extreme population feminisation and collapse predicted by the end of the century (Lockley and Eizaguirre, 2021). To gain molecular insights into this mechanism, we compared the chromosomal locations of 199 TSD-linked genes present in our loggerhead genome, VGP green, and VGP leatherback genomes. Consistent with Bentley et al. (2023), most TSD-linked genes are single-copy and reside on equivalent chromosomes in sea turtle species (**Figure 5A, 5B**). Only the EP300 (E1A binding protein p300) gene was not syntenic between loggerheads and greens (**Figure 5A**). Although it mapped to chromosome 1 in all species, a duplicate on chromosome 10 was found in the green and leatherback genomes. Validation is required to resolve if this represents a true duplicate in greens and leatherbacks that has been lost in loggerheads. Two genes were syntenic between loggerheads and greens but not leatherbacks (**Figure 5B**): CIRBP (Cold-inducible RNA-binding protein; loggerhead/green: chromosome 25; leatherback: chromosome 27) and IFIT5 (Interferon Induced Protein with Tetratricopeptide Repeats 5; loggerhead/green: chromosome 7; leatherback: chromosome 1). These may reflect fission or fusion events after the divergence of the Cheloniidae and Dermochelyidae lineages. As rearrangements and copy number variants can alter gene regulation (Harewood and Fraser, 2014), those three genes may contribute to inter-species differences in the TSD response curve and should be investigated. Nevertheless, such structural variants seem rare in sea turtles overall, unlike in other Testudines (Valenzuela and Adams, 2011; Lee *et al*., 2019). Finer scale genetic variation may instead be more important, particularly within species. For example, a SNP on the CIRBP gene, which we found to be non-syntenic against leatherbacks, influences TSD in snapping turtles (*Chelydra serpentina*) and exhibits varying allele frequencies with latitude (Schroeder *et al*., 2016). It is thus crucial to continue characterising genetic variants that contribute to the adaptive potential of the TSD mechanism, to evaluate whether different sea turtle species and populations can evolve to mitigate sex ratio skew in the face of rising temperatures as climate change progresses.

**Figure 5.**
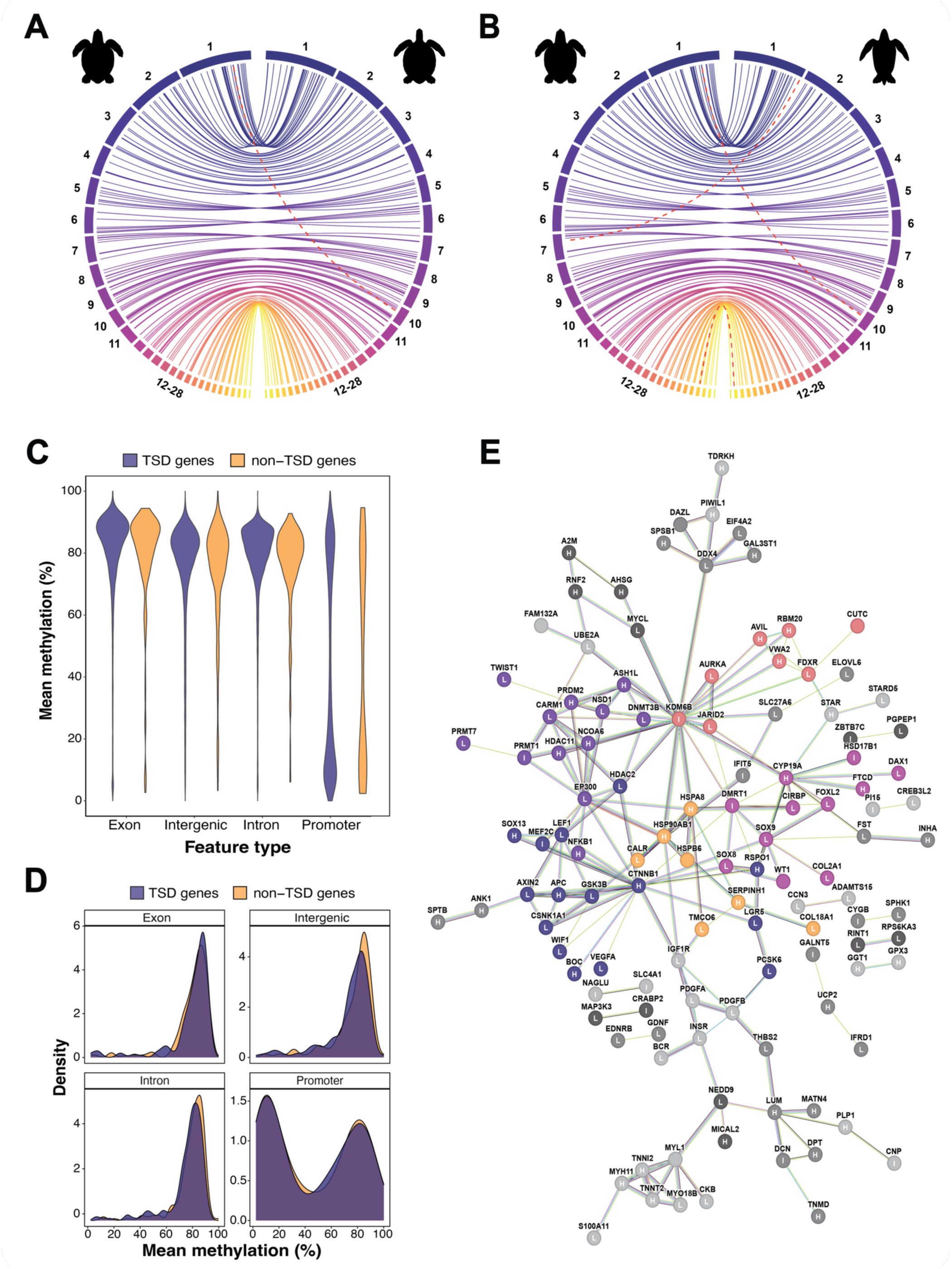
Molecular insights into TSD-linked genes: synteny, methylation and functional associations. (A-B) Chromosomal locations of 199 TSD-linked genes across 205 loci in our loggerhead assembly against the Vertebrate Genomes Project **(A)** green assembly and **(B)** leatherback assembly. In both Circos plots, chromosomes of the loggerhead assembly are plotted on the left, with colours representing the 28 chromosomes. Dashed red lines indicate genes that mapped to different locations between species (loggerhead versus green: EP300, loggerhead versus leatherback: EP300, CIRBP, IFIT5). There is overall high synteny of TSD- linked genes between sea turtle species. (**C)** Violin plots of mean methylation (%) per gene by feature type for TSD-linked (n=200 genes; purple) versus non-TSD-linked genes (n=15,041 single-copy orthologues between sea turtle species; yellow). **(D)** Density plots of mean methylation (%) per gene by feature type between 200 TSD-linked genes (purple) and a random subset of 200 non-TSD-linked genes (yellow). Both plots show that methylation is distributed differently across feature types, but similarly between TSD-linked versus non-TSD-linked genes. **(E)** Predicted functional association network of proteins coded by TSD-linked genes, created with STRING database. 119 genes with at least one connection are shown (n=28 clusters total). Nodes represent gene IDs and edges represent associations. Edge colours represent different evidence types used for association prediction. Non-grey node colours represent the five largest functional clusters identified via Markov Clustering. Letters on nodes represent methylation status of gene promoters in the reference individual methylome. Categories are based on the bimodal methylation distribution observed in **Figure 5D**: H: highly methylated (>70%), L: lowly methylated (<30%), I: intermediate methylation (30-70%). 67 promoters (33.8%) were hyper-methylated, 96 (48%) were hypo-methylated and 36 (18.2%) had intermediate methylation. Proportions of each promoter methylation category were different amongst the five largest functional clusters (χ^2^=41.697, p<0.0001).

#### TSD-linked genes: methylation patterns

Given genetic differences between sea turtles are low and epigenetic regulation can contribute to the adaptive potential of TSD species (Piferrer, 2021; Balard *et al*., 2024), we next provide insights into the methylation status of TSD-linked genes. We tested if methylation differed between 200 TSD-linked genes present in the loggerhead genome, versus 1000 random subsets of 200 non-TSD-linked genes sampled from 15,041 single-copy orthologues identified between sea turtle species. Mean methylation was not associated to gene category (TSD-linked or non-TSD-linked genes), as supported by only 155 out of 1000 (15.5%) tests crossing a ‘significance threshold’ of p<0.05 (**Figure S7**). The interaction between gene category and feature type did not reveal differences either, with 38 out of 1000 (0.38%) tests passing a threshold of p<0.05 (**Figure S7**), and similar methylation distributions observed across all feature types (**Figure 5C, 5D**). Instead, feature type was the sole determinant of mean methylation, with all 1000 tests (100%) passing p<0.05 (**Figure S7**), as expected given the importance of genomic context on methylation patterns and functions (Jones, 2012). Similar results were observed when testing associations against the proportion of highly methylated CpGs (**Figure S8**). Exons, introns and gene-associated intergenic space (<10 kbp from TSS) were mostly hypermethylated, with a peak of methylation centred on ∼80% (**Figure 5C, 5D**). In contrast, methylation was bimodally distributed for promoters, with peaks of low (∼10%) and high (∼80%) methylation (**Figure 5C, 5D**). These distributions are consistent with previously described patterns in vertebrates (Elango and Yi, 2008; Keller, Han and Yi, 2016).

We characterised large-scale methylation patterns from blood. This is a valuable tissue for conservation monitoring as it can be collected via minimally invasive sampling, yet still report on an individual’s physiology (De Paoli-Iseppi *et al*., 2017; Viitaniemi *et al*., 2019; Grant *et al*., 2022), health (Yousefi *et al*., 2022) and environmental exposure (Xu *et al*., 2020; Mäkinen *et al*., 2022). For example, a non-lethal method for sexing sea turtle hatchlings is urgently required to assess nest sex ratios as global temperatures continue to rise (Lockley and Eizaguirre, 2021). Sixteen differentially methylated regions were identified from skin samples of adults but not earlier life-stages of green turtles via reduced representation sequencing (Mayne *et al*., 2023). Methylome-wide discovery scans should thus be used to identify biomarkers of hatchling sex from blood samples, as conducted in relation to thermal stress in a study that employed this reference assembly (Yen *et al*., 2023).

#### TSD-linked genes: a functional association map

Using the STRING database (Szklarczyk *et al*., 2020), we built a network of functional associations between proteins coded by 191 TSD-linked genes **(Figure 5E**). In total, 119 TSD- linked genes had at least one connection identified via Markov Clustering (n=28 clusters total), and the five largest clusters encompassed 54 TSD-linked genes (45.4%). Members of the biggest cluster (n=15, dark blue) were mainly linked to Wnt signalling. Connections were centred on CTNNB1 (beta-catenin), which is critical for female determination in vertebrates (Mork and Capel, 2013; Liu *et al*., 2022). The next largest cluster (n=12, dark purple) was primarily composed of epigenetic regulators. The most connected protein was EP300 (E1A binding protein p300), a histone acetyltransferase that controls cell proliferation, with germ cell numbers proposed to influence TSD in a freshwater turtle (Tezak *et al*., 2023). The third cluster (n=11, mauve) contained genes with well-described roles in TSD and steroid hormone signalling (Rhen and Schroeder, 2010). SOX9 (SRY-related HMG box gene 9), a key regulator of male differentiation (Kent *et al*., 1996; Moreno-Mendoza, Harley and Merchant-Larios, 2001), was most connected. Although the functional nature of the fourth cluster (n=8, pink) was less obvious, the most connected protein was the histone demethylase KDM6B (Lysine Demethylase 6B), demethylase which is causally linked to temperature-dependent male determination in freshwater turtles (Ge *et al*., 2018). The final cluster contains heat shock proteins and chaperones (n=7, yellow), which could play a role in temperature sensing and transduction during TSD (Kohno *et al*., 2009). These identified clusters already reveal patterns surrounding TSD in sea turtles. Sensory systems associated with heat shock responses could act as environmental messengers that link with epigenetic regulators to alter gene expression, cell proliferation, and subsequently activate known endocrine systems. With growing genomic tools for sea turtles, this hypothesis will need to be tested.

Given the prevalence of epigenetic regulators in the functional association network, we overlaid promoter methylation status of TSD-linked genes from our loggerhead blood methylome (**Figure 5E**). In total, 67 genes (33.8%) were highly (>70%) methylated, 96 genes (48.0%) were lowly (<30%) methylated, and 36 genes (18.2%) had intermediate methylation.

The proportions of each promoter methylation category differed amongst the five largest functional clusters (χ^2^=41.697, p<0.0001, **Table S9**). The lowest methylated clusters involved Wnt signalling genes (dark blue; high: 33.3%, intermediate: 6.7%, low: 60%) and the well-described TSD-linked/hormone signalling genes (mauve; high: 20%, intermediate: 20%, low: 60%). No clusters were hyper-methylated, as the heat shock cluster (yellow) contained the largest proportion of highly methylated genes, with an equal split between highly and lowly methylated genes (high: 50%, intermediate: 0%, low: 50%). With promoter methylation classically linked to transcriptional repression (Jones, 2012), we may have captured female-specific methylation signatures on a gene network scale. By integrating methylation and functional association information into a single map, we facilitate the investigation into the relationships between gene function, connectivity, and epigenetic regulation in the context of TSD for sea turtles.

## CONCLUSIONS

To support conservation efforts of threatened loggerhead sea turtles, as well as all sea turtles and TSD species in general, we present a chromosome-scale genome representing the globally significant East Atlantic nesting group. Our reference assembly will enhance genomic inferences for this population, and contribute a new genome for comparative studies within and between species. Moreover, we anticipate our ONT-derived methylome can guide future epigenetic study design. To demonstrate their potential, we applied these novel resources to yield a variety of insights relevant to the conservation of sea turtles. At the population level, we emphasise the role of climate change and niche availability on effective population size fluctuations. On the chromosome level, we recommend microchromosomes as special regions for monitoring functional genetic and epigenetic diversity. Finally, we bring gene-level insights into the TSD cascade of sea turtles, highlighting three TSD-linked genes with potential inter-species rearrangements, and providing a map of functional associations and methylation status for TSD-linked genes to assist sex biomarker discovery. By simultaneously generating genome and methylome resources to provide molecular insights in loggerhead sea turtles, our study showcases the application of this dual framework for informing the conservation of an endangered, flagship species.

## DATA AVAILABILITY

All genomic resources and sequencing data reported in this article are being uploaded to ENA (European Nucleotide Archive) under study accession PRJEB79015. All supporting data and resources will be made available. All scripts are being deposited to the GitHub repository: https://github.com/eugeniecyen/Article_CarCar_GenomeAssembly.

## LIST OF ABBREVIATIONS

5mC: 5-methylcytosine, 5hmC: 5-hydroxymethylcytosine, bp: base pairs, CBP: Canada BioGenome Project, CpG: 5’-C-phosphate-G-3, ENA: European Nucleotide Archive, Gbp: giga base pairs, kbp: kilo base pairs, Mbp: mega base pairs, Mya: million years ago, Ne: effective population size, ONT: Oxford Nanopore Technology, PCR: polymerase chain reaction, PSMC: Pairwise Sequentially Markovian Coalescent, TSS: transcription start site, VGP: Vertebrate Genomes Project, WGBS: whole genome bisulfite sequencing

## SUPPLEMENTARY FILES

**Table S1.** Publicly available RNA-Seq reads mined for genome annotation

**Table S2.** Metadata for ten nesting loggerheads sampled for WGBS

**Table S3.** WGBS summary statistics for ten nesting loggerheads

**Table S4.** Full BUSCO summary for genome assemblies across sea turtle species

**Table S5.** Repetitive element summary statistics

**Table S6.** Genome annotation summary statistics

**Table S7.** Full BUSCO summary for genome annotations across sea turtle species

**Table S8.** Summary of genome alignment identity between sea turtle species

**Table S9.** Proportions of promoter methylation categories in the top five functional clusters

**Text S1.** Extended methods and results for mitochondrial genome assembly and annotation

**Text S2.** Extended methods for methylation calling from WGBS data of ten loggerheads

**Text S3.** Extended methods for annotating gene feature types

**Text S4.** Extended methods for identification and curation of TSD-linked genes

Figure S1. GenomeScope profile for our loggerhead genome

Figure S2. Auxiliary PSMC tests with 100 bootstraps

Figure S3. GC-coverage blob-plot to evaluate assembly contamination

Figure S4. Circos plot of our loggerhead mitochondrial assembly and annotation

Figure S5. K-mer spectrum plot of our loggerhead assembly

Figure S6. Comparison of genome properties between macro- and microchromosomes

Figure S7. P-value histogram from comparing mean methylation between TSD-linked genes and non-TSD-linked genes

Figure S8. P-value histogram from comparing the proportion of highly methylated CpGs between TSD-linked genes and non-TSD-linked genes

## DECLARATIONS

### Ethics approval

All sample collection adhered to national legislation and were approved by the Direção Nacional do Ambiente de Cabo Verde (Permits: 13/DNA/2020, 037/DNA/2021).

### Author contributions

E.C.Y and C.E designed the study. E.C.Y, C.E, A.T and K.F collected blood samples. E.C.Y performed genome assembly, annotation and quality assessment, with contributions from J.M.M-D. A.B, E.C.Y and D-M.J.T performed DNA methylation analysis of ONT reads. E.C.Y performed sample processing and methylation analysis for WGBS. E.C.Y performed genome property and synteny analyses. E.C.Y performed PSMC, with input and guidance on geoclimatic events from H.L.F. J.D.G identified TSD-linked genes, orthogroup genes, mapped chromosomal locations and produced the functional association network. E.C.Y analysed methylation of TSD-linked genes. E.C.Y wrote the manuscript with contributions from C.E, J.D.G, H.L.F, S.J.R, J.M.M-D and feedback from all authors.

## Acknowledgements

The authors thank all staff with Project Biodiversity (Sal, Cabo Verde) for their support in the field. We also thank Chloe Economou (Queen Mary University of London, UK) for conducting DNA extractions for the reference individual and William Tyne (Queen Mary University of London, UK) for ONT library preparation and sequencing.

