## Supplementary Material for "Chromosome-level genome assembly and methylome profile enables insights for the conservation of endangered loggerhead sea turtles"

| BioProject | SRA Run | BioSample | Associated publication | Tissue | Life stage | Sex | Treatment | Raw bases (Gbp) | Raw reads | Passed reads | Hint type generated |
|---|---|---|---|---|---|---|---|---|---|---|---|
| PRJNA663187 | SRR12630873 | SAMN16122281 | Chow et al. (2021) | Gonad | Hatchling | Female | None | 10.2 | 50746307 | 49954134 (98.44%) | Transcript |
| PRJNA663187 | SRR12630874 | SAMN16122280 | Chow et al. (2021) | Gonad | Hatchling | Female | None | 10.3 | 51399125 | 50152916 (97.58%) | Transcript |
| PRJNA663187 | SRR12630879 | SAMN16122282 | Chow et al. (2021) | Gonad | Hatchling | Male | None | 9.3 | 46557205 | 45820268 (98.42%) | Transcript, transcriptome |
| PRJNA663187 | SRR12630881 | SAMN16122283 | Chow et al. (2021) | Gonad | Hatchling | Male | None | 2.2 | 11018312 | 10879472 (98.74%) | Transcript |
| PRJNA560561 | SRR10032986 | SAMN12591173 | Hernández-Fernández et al. (2021) | Blood | Hatchling | Unknown | None | 15.7 | 77607764 | 76009322 (97.94%) | Transcript |
| PRJNA560561 | SRR10032987 | SAMN12591166 | Hernández-Fernández et al. (2021) | Blood | Hatchling | Unknown | None | 16.2 | 80140038 | 78357616 (97.78%) | Transcript |
| PRJNA560561 | SRR10032988 | SAMN12591165 | Hernández-Fernández et al. (2021) | Blood | Hatchling | Unknown | None | 16.4 | 81364511 | 79461565 (97.66%) | Transcript |
| PRJNA560561 | SRR10032989 | SAMN12591164 | Hernández-Fernández et al. (2021) | Blood | Adult | Female | None | 5.8 | 28500836 | 27552342 (96.67%) | Transcript |
| PRJNA560561 | SRR10032990 | SAMN12591163 | Hernández-Fernández et al. (2021) | Blood | Adult | Male | None | 5.6 | 27498122 | 26591514 (96.70%) | Transcript |
| PRJNA560561 | SRR10032991 | SAMN12591162 | Hernández-Fernández et al. (2021) | Blood | Juvenile | Female | None | 5 | 24727270 | 23791771 (96.22%) | Transcript |
| PRJNA560561 | SRR10032992 | SAMN12591161 | Hernández-Fernández et al. (2021) | Blood | Juvenile | Female | None | 5.4 | 26568011 | 25725953 (96.83%) | Transcript |
| PRJNA560561 | SRR10032993 | SAMN06350885 | Hernández-Fernández et al. (2021) | Blood | Juvenile | Male | None | 5.4 | 26826849 | 25946591 (96.72%) | Transcript |
| PRJNA649079 | SRR12335440 | SAMN15657957 | NA | Brain | Hatchling | Unknown | None | 3.2 | 10443220 | 9989577 (95.66%) | Transcript, transcriptome |
| PRJNA649079 | SRR12335451 | SAMN15657956 | NA | Heart | Hatchling | Unknown | None | 2.2 | 7281215 | 6917002 (95.00%) | Transcript, transcriptome |
| PRJNA649079 | SRR12335462 | SAMN15657955 | NA | Brain | Hatchling | Unknown | None | 10.7 | 35334430 | 33529137 (94.89%) | Transcript |
| PRJNA649079 | SRR12335473 | SAMN15657962 | NA | Heart | Hatchling | Unknown | Heat shock | 6 | 19906468 | 19075205 (95.82%) | Transcript |
| PRJNA649079 | SRR12335484 | SAMN15657961 | NA | Brain | Hatchling | Unknown | Heat shock | 5.8 | 19324895 | 18544037 (95.96%) | Transcript |
| PRJNA649079 | SRR12335506 | SAMN15657959 | NA | Brain | Hatchling | Unknown | Heat shock | 3.8 | 12717687 | 12169458 (95.69%) | Transcript |
| PRJNA649079 | SRR12335527 | SAMN15657964 | NA | Heart | Hatchling | Unknown | Heat shock | 2.6 | 8671833 | 8296669 (95.67%) | Transcript |
| PRJNA649079 | SRR12335528 | SAMN15657963 | NA | Brain | Hatchling | Unknown | Heat shock | 4.1 | 13668130 | 13062514 (95.57%) | Transcript |
| PRJNA649079 | SRR12335529 | SAMN15657954 | NA | Heart | Hatchling | Unknown | None | 1.7 | 5736034 | 5441041 (94.86%) | Transcript |
| PRJNA649079 | SRR12335530 | SAMN15657953 | NA | Brain | Hatchling | Unknown | None | 7.7 | 25603564 | 24423753 (95.39%) | Transcript |
| PRJNA339812 | SRR5330501 | SAMN06350885 | Hernández-Fernández et al. (2017) | Blood | Juvenile | Female | None | 5.4 | 26826849 | 25998957 (96.91%) | Transcript |
| PRJNA660024 | SRR12540984 | SAMN15933003 | Banerjee et al. (2021) | Blood | Unknown | Unknown | None | 8.3 | 27664060 | 23059350 (83.35%) | Transcript |

**Table S1. Publicly available RNA-Seq reads mined for genome annotation.** Reads were downloaded from the NCBI Sequence Read Archive (Leinonen, Sugawara and Shumway, 2011). These comprised of 746,132,735 reads from 24 loggerheads across three life stages (hatchling, juvenile and adult), four tissue types (blood, gonad, brain and heart) and both sexes.

| Sample ID | Date | Locality | Island | Country | Latitude | Longitude |
|---|---|---|---|---|---|---|
| SLL063 | 29/07/2021 | Algodoeiro Beach | Sal | Cabo Verde | 16.62028°N | -22.92938°E |
| SLL065 | 29/07/2021 | Algodoeiro Beach | Sal | Cabo Verde | 16.62042°N | -22.92945°E |
| SLL142 | 29/07/2021 | Algodoeiro Beach | Sal | Cabo Verde | 16.61960°N | -22.92927°E |
| SLL143 | 29/07/2021 | Algodoeiro Beach | Sal | Cabo Verde | 16.61625°N | -22.92858°E |
| SLL144 | 29/07/2021 | Algodoeiro Beach | Sal | Cabo Verde | 16.61834°N | -22.55732°E |
| SLL146 | 29/07/2021 | Algodoeiro Beach | Sal | Cabo Verde | 16.62004°N | -22.92929°E |
| SLL171 | 29/07/2021 | Algodoeiro Beach | Sal | Cabo Verde | 16.61585°N | -22.92860°E |
| SLL176 | 29/07/2021 | Algodoeiro Beach | Sal | Cabo Verde | 16.61627°N | -22.92860°E |
| SLL188 | 29/07/2021 | Algodoeiro Beach | Sal | Cabo Verde | 16.61597°N | -22.92858°E |
| SLL189 | 29/07/2021 | Algodoeiro Beach | Sal | Cabo Verde | 16.61633°N | -22.92857°E |

**Table S2. Metadata for ten nesting loggerheads sampled for WGBS.**

| Sample ID | Total read pairs | Mean mapping efficiency (%) | Bisulfite conversion efficiency (%) | Total CpGs after de-stranding | Mean CpG coverage after de-stranding |
|---|---|---|---|---|---|
| SLL063 | 132,232,345 | 79.0 | 99.30 | 25,445,337 | 9.07 |
| SLL065 | 132,185,458 | 82.2 | 99.38 | 25,509,991 | 9.58 |
| SLL142 | 132,204,488 | 82.6 | 99.39 | 25,492,037 | 9.36 |
| SLL143 | 132,161,944 | 84.3 | 99.37 | 25,519,785 | 9.70 |
| SLL144 | 132,219,979 | 81.6 | 99.38 | 25,499,082 | 9.47 |
| SLL146 | 132,165,428 | 77.0 | 99.36 | 25,501,839 | 9.20 |
| SLL171 | 132,186,272 | 78.5 | 99.47 | 25,417,602 | 8.90 |
| SLL176 | 132,158,064 | 76.0 | 99.43 | 25,501,178 | 8.99 |
| SLL188 | 132,272,286 | 75.8 | 99.43 | 25,519,119 | 9.16 |
| SLL189 | 132,138,954 | 74.4 | 99.43 | 25,449,602 | 8.63 |

**Table S3. WGBS summary statistics for ten nesting loggerheads.** Alignment, de-duplication and methylation calling was performed with Bismark v.0.22.1 (Krueger and Andrews, 2011).

| Assembly | Complete BUSCOs (%) | Single copy BUSCOs (%) | Duplicated BUSCOs (%) | Fragmented BUSCOs (%) | Missing BUSCOs (%) |
|---|---|---|---|---|---|
| **CarCar_QM_v1_2021_12_Scaff** (Loggerhead turtle, scaffolded) | 97.1 | 96.2 | 0.9 | 0.4 | 2.5 |
| **CarCar_GSC_CCare_1.0** (Loggerhead turtle, scaffolded) | 96.1 | 95.2 | 0.9 | 0.4 | 3.5 |
| **rDerCor1.pri.v4** (Leatherback turtle, scaffolded) | 96.3 | 95.3 | 1.0 | 0.6 | 3.1 |
| **rCheMyd1.pri.v2** (Green turtle, scaffolded) | 97.2 | 96.2 | 1.0 | 0.4 | 2.4 |
| **CheMyd_1.0** (Green turtle, contig-level) | 95.9 | 94.8 | 1.1 | 1.3 | 2.8 |

**Table S4. Full BUSCO summary for genome assemblies across sea turtle species.** BUSCO scores were calculated against the Sauropsida gene set (n=7480) with BUSCO v.5.1.2 in genome mode (Simão *et al.*, 2015).

| Element type | Total count | Total length (bp) | % of sequence |
|---|---|---|---|
| **Retroelements** | 2,274,746 | 588,778,218 | 27.4 |
| SINEs | 304,426 | 30,730,756 | 1.43 |
| LINEs | 694,162 | 274,850,101 | 12.8 |
| LTRs | 1,276,158 | 283,197,361 | 13.2 |
| **DNA transposons** | 1,439,663 | 224,774,317 | 10.5 |
| **Unclassified** | 697,775 | 100,371,114 | 4.68 |

**Table S5. Repetitive element summary statistics.** Statistics generated by RepeatMasker v.4.1.4 (Smit, Hubley, and Green 2022).

| Annotation statistics | |
|---|---|
| Total repeat sequence masked (Mbp) | 924.6 (43.1% of genome) |
| Total genes | 29,883 |
| Total genes with functional annotations | 23,690 (82.0% of genes) |
| Total genes with GO annotations | 16,954 (58.7% of genes) |
| Total exons | 272,673 |
| Total introns in coding sequence | 241,823 |
| Mean gene length (kbp) | 40.1 |
| Mean exon length (bp) | 170 |
| Mean intron length in coding sequence (bp) | 4799 |

**Table S6. Genome annotation summary statistics.** Annotation statistics were computed with AGAT v.0.9.1 (Dainat, 2022).

| Assembly | Complete BUSCOs (%) | Single copy BUSCOs (%) | Duplicated BUSCOs (%) | Fragmented BUSCOs (%) | Missing BUSCOs (%) |
|---|---|---|---|---|---|
| **CarCar_QM_v1_2021_12_Scaff** (Loggerhead turtle, scaffolded) | 94.7 | 93.9 | 0.8 | 1.1 | 4.2 |
| **CarCar_GSC_CCare_1.0** (Loggerhead turtle, scaffolded) | 97.8 | 96.8 | 1.0 | 0.2 | 2.0 |
| **rDerCor1.pri.v4** (Leatherback turtle, scaffolded) | 98.2 | 97.1 | 1.1 | 0.5 | 1.3 |
| **rCheMyd1.pri.v2** (Green turtle, scaffolded) | 98.9 | 97.8 | 1.1 | 0.3 | 0.8 |
| **CheMyd_1.0** (Green turtle, contig-level) | 95.8 | 94.9 | 0.9 | 1.5 | 2.7 |

**Table S7. Full BUSCO summary for genome annotations across sea turtle species.** BUSCO scores were calculated on the longest gene isoforms against the Sauropsida gene set (n=7480) with BUSCO v.5.1.2 in protein mode (Simão *et al.*, 2015).

| Alignment identity | VGP green turtle | VGP leatherback turtle |
|---|---|---|
| No match | 2.81 | 7.57 |
| <25% | 0.10 | 1.47 |
| 25-50% | 7.01 | 87.6 |
| 50-75% | 90.1 | 3.37 |
| >75% | 0.01 | 0.00 |

**Table S8. Summary of genome alignment identity between sea turtle species.** Alignments and statistics produced by D-GENIES v.1.5.0 (Cabanettes and Klopp, 2018)

| Mean promoter methylation status | Proportion (%) per cluster | | | | |
|---|---|---|---|---|---|
| | Dark blue | Dark purple | Mauve | Pink | Yellow |
| High | 33.33 | 41.67 | 20 | 42.86 | 50 |
| Intermediate | 6.67 | 8.33 | 20 | 14.29 | 0 |
| Low | 60.00 | 50.00 | 60 | 42.86 | 50 |

**Table S9. Proportions of promoter methylation categories in the top five functional clusters.** Methylation status of TSD-linked gene promoters in the reference individual methylome (i.e. blood of a nesting female) were assigned into high (>70%), intermediate (30-70%) or low (<30%) categories based on the bimodal distribution observed (**Figure 5D**). The proportion of gene promoters falling into these three categories are provided for the five largest functional clusters identified via Markov Clustering from our STRING functional association network created for TSD-linked genes (**Figure 5E**).

**Text S1. Extended methods and results for mitochondrial genome assembly and annotation.** We assembled and annotated the mitochondrial genome from our Illumina data. To extract a read set enriched for mitochondrial sequence, reads were mapped via BWA-MEM v.0.7.17 (Li, 2013) against a published loggerhead mitochondrial assembly (Drosopoulou et al., 2012). Mapped reads were inputted to MitoZ v.3.4 (Meng et al., 2019) with the Megahit assembler. Our assembly consisted of a circular 16,574 bp contig with 37 genes (13 protein coding genes, 22 tRNA genes and 2 rRNA genes, **Figure S4**). This assembly was 98.9% identical to a loggerhead mitochondrial genome from a Greek population (Drosopoulou et al., 2012). Divergence likely reflects genetic structure between the North Atlantic and Mediterranean nesting groups, driven by female philopatry in sea turtles (Baltazar-Soares et al., 2020; Tolve et al., 2023). To assign the reference individual to a mitochondrial haplogroup, control region haplotype sequences found in the Cabo Verde nesting group (Baltazar-Soares et al., 2020) were downloaded from NCBI GenBank (Sayers et al., 2021) and aligned against our assembly via BLASTn v.2.7.1+ (Altschul et al., 1990). The control region was 99.9% similar to the CC-A1.4 haplotype (GenBank ID: EU179439.1). The reference individual is therefore a member of Haplogroup I (CC-A1), the oldest and commonest loggerhead lineage in Cabo Verde (Baltazar-Soares et al., 2020).

**Text S2. Extended methods for methylation calling from WGBS data of ten loggerheads.** Raw WGBS reads were trimmed for remaining adapters and filtered for a mean Phred score >Q20 using cutadapt v.2.10 (Martin, 2011). Using Bismark v.0.22.1 with default options (Krueger and Andrews, 2011), trimmed reads were aligned against our reference assembly, giving a mean mapping efficiency of 79.1 ± 3.37 (SD) % per sample (**Table S3**). Alignments were deduplicated with Bismark in paired end mode, followed by merging and sorting with samtools v.1.9 (Li *et al.*, 2009). Methylation calling was then performed in Bismark. Percentage methylation totalled across CHG and CHH sites were calculated to obtain an estimate of bisulfite conversion efficiency (Laine *et al.*, 2023). This gave a mean of 99.39 ± 0.047 (SD) %, indicating high conversion efficiency (**Table S3**). To improve coverage and minimise pseudo-replication, we further de-stranded adjacent cytosines per CpG site using the 'merge_CpG.py' script (Cristofari, 2023), as methylation occurs symmetrically at CpG sites in vertebrates (Klughammer *et al.*, 2023). On average, this resulted in 25,485,557 ± 35,209 (SD) CpG sites per sample, with a de-stranded coverage of 9.2 ± 0.33 (SD) X (**Table S3**). Methylation calls at CpG sites were further processed in RStudio v.4.2.2 (R Core Team, 2021) with the methylKit package v.1.24.0 (Akalin *et al.*, 2012). CpG sites were excluded if they had a coverage lower than 5X, or if they were within the 99.9th percentile to account for possible PCR bias (Wreczycka *et al.*, 2017). Coverage was then normalised between samples using methylKit's 'normalizeCoverage' function. As a final filtering step, we retained CpG sites that were covered in at least 75% of individuals, leaving 24,299,151 CpG sites for downstream analyses. For each CpG site, percentage methylation was calculated per individual with methylKit's 'percMethylation' function, then the mean across all individuals was calculated per CpG site to provide an average WGBS methylome representative of the population.

**Text S3. Extended methods for annotating gene feature types.**

To annotate the feature type upon which CpG sites reside, we used the R packages genomation v.1.30.0 (Akalin *et al.*, 2015) and GenomicRanges v.1.50.2 (Lawrence *et al.*, 2013) alongside our reference genome. Promoter regions were defined as 1500 bp upstream and up to 500 bp downstream of a transcriptional start site (TSS; Heckwolf *et al.*, 2020). All CpG sites were assigned to one of four feature types using genomation's 'annotateWithGeneParts' function, in the following order of precedence when features overlapped: promoter, exon, intron or intergenic region. To attach functional gene information to our DMS, those in genic regions (i.e., located on a promoter, intron or exon) were associated to a gene using the 'findOverlaps' function of GenomicRanges. DMS in intergenic regions were associated to a gene using genomation's 'getAssociationWithTSS' function, if they were less than 10 kb away from the nearest TSS (Heckwolf *et al.*, 2020).

**Text S4. Extended methods for identification and curation of TSD-linked genes.**

Bentley et al. (2023) compiled the most comprehensive list to date of 223 genes with documented links to temperature-dependent sex determination (TSD) pathways. Firstly, we manually curated this list by removing 11 genes that were not found in either the green or leatherback genomes, consolidating GLU and NAGLU which referred to the same gene, and correcting the gene name 'ST6GALC2' to 'ST6GAL2'. We further replaced four sequences that did not map to the corresponding gene: the A2M sequence was replaced to XM_037912788.2 for the green turtle, the PDGFA sequence was replaced to XR_003565502.3 in the green turtle and XR_005295469.2 in the leatherback turtle, the PDGFB sequence was replaced to XM_037881931.2 in the green turtle and XM_038387707.2 in the leatherback turtle, and ST6GAL2 was replaced to XM_007054611.4 in the green turtle and XM_043509904.1 in the leatherback turtle.

Using our curated list of TSD-linked genes, orthologues were identified in our loggerhead assembly via BLASTn v.2.11.0 with parameters '-evalue 1e-30' and '-perc_identity 70' (Altschul *et al.*, 1990) against gene sequences from the VGP green turtle annotation, which is more closely related to the loggerhead turtle than the leatherback turtle. To verify that true orthologues were selected, a manual curation step integrating BLAST homology and gene name information was conducted. This involved looking up the gene name for every BLAST hit in our loggerhead functional annotation, and retaining those that matched the query gene name. We took the top BLAST hit if there was no match to query gene name, due to genes having multiple aliases or missing annotation information. For loggerhead genes that matched sequences in two chromosomal locations, hits were retained if they were both syntenic or neither syntenic with the other species. If one sequence was syntenic and the other was not, the non-syntenic sequence was removed under the conservative assumption of it being an assembly or orthologue identification error. This left 201 unique TSD-linked genes in our loggerhead genome for downstream analyses (five genes annotated in two locations; 206 total loci).

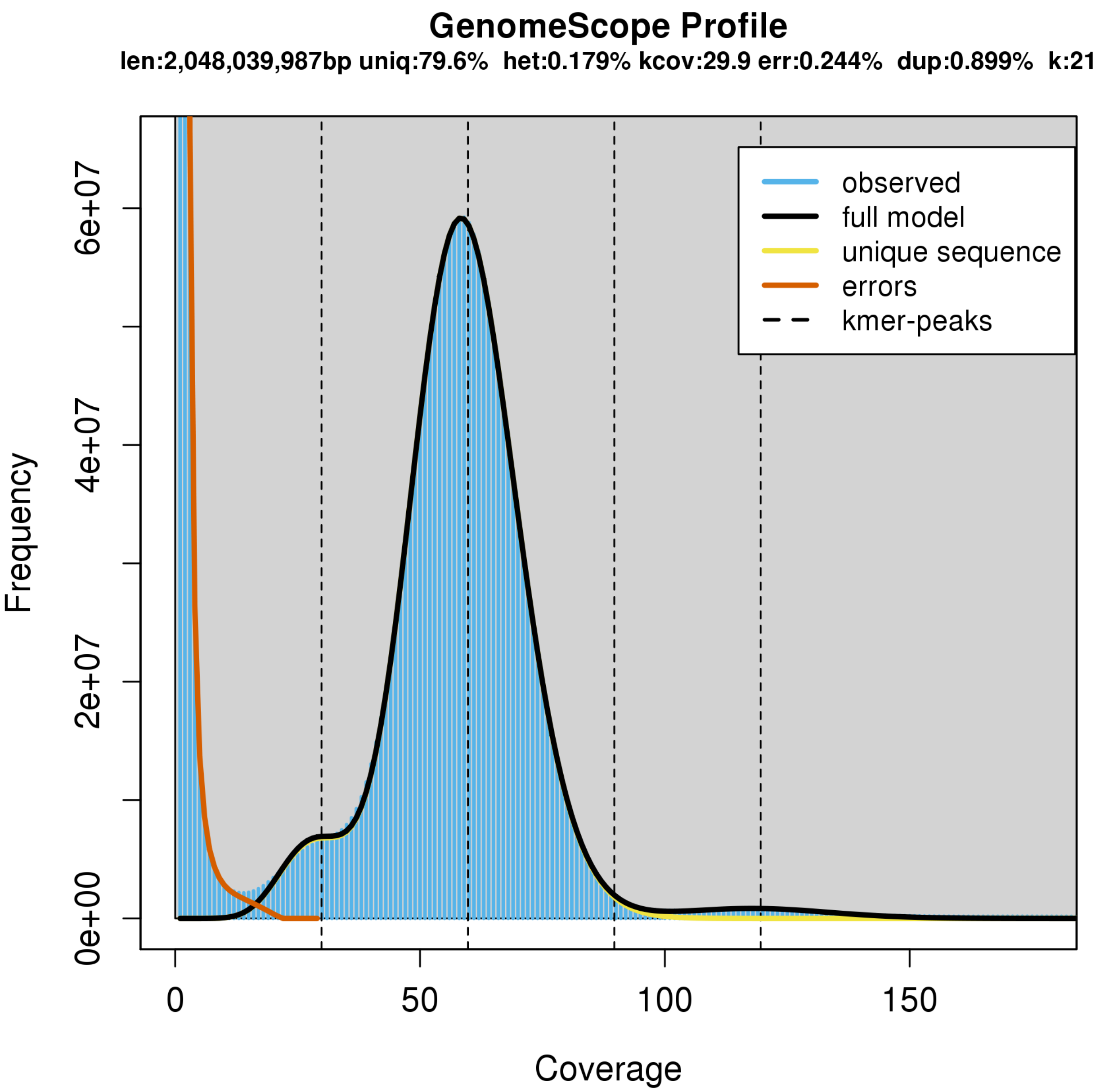


**Figure S1. GenomeScope profile for our loggerhead genome.** Produced using GenomeScope (Vurture *et al.*, 2017) on k-mers derived from our raw, Illumina reads, with parameter k=21. Estimated haploid genome size is 2.05 Gbp, heterozygosity is 0.179% and repeat fraction is 20.4%. These metrics were used to help choose downstream parameters.

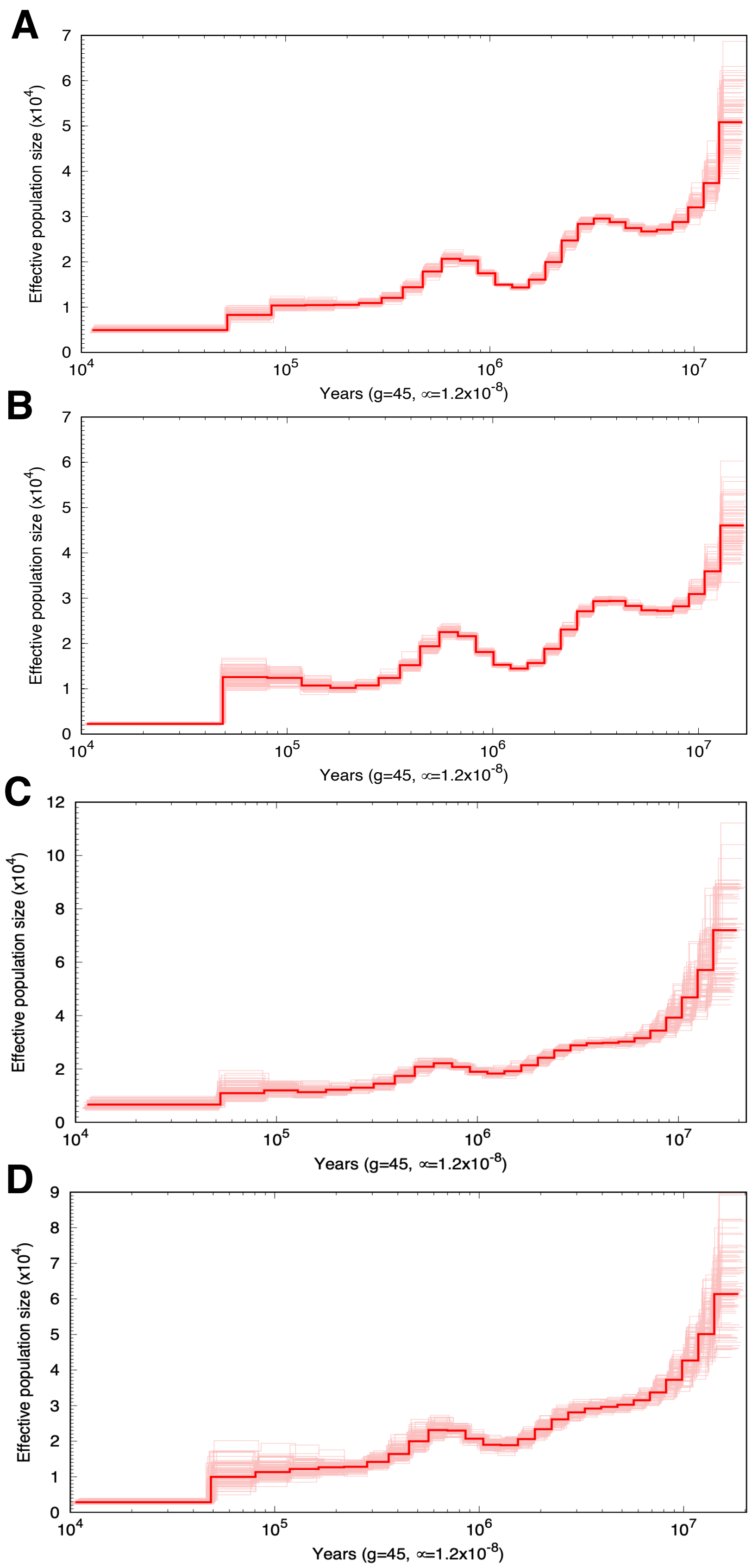


**Figure S2. Auxiliary PSMC tests with 100 bootstraps**. PSMC run from the 11 macrochromosomes of the loggerhead genome (80.8% of assembly) for **(A)** SLK063 (Cabo Verde, East Atlantic), and **(B)** SAMN20502673 (Brazil, West Atlantic). PSMC was also run on the 17 microchromosomes (19.2% of assembly) for **(C)** SLK063 and **(D)** SAMN20502673, confirming overall patterns were comparable with macrochromosomes.

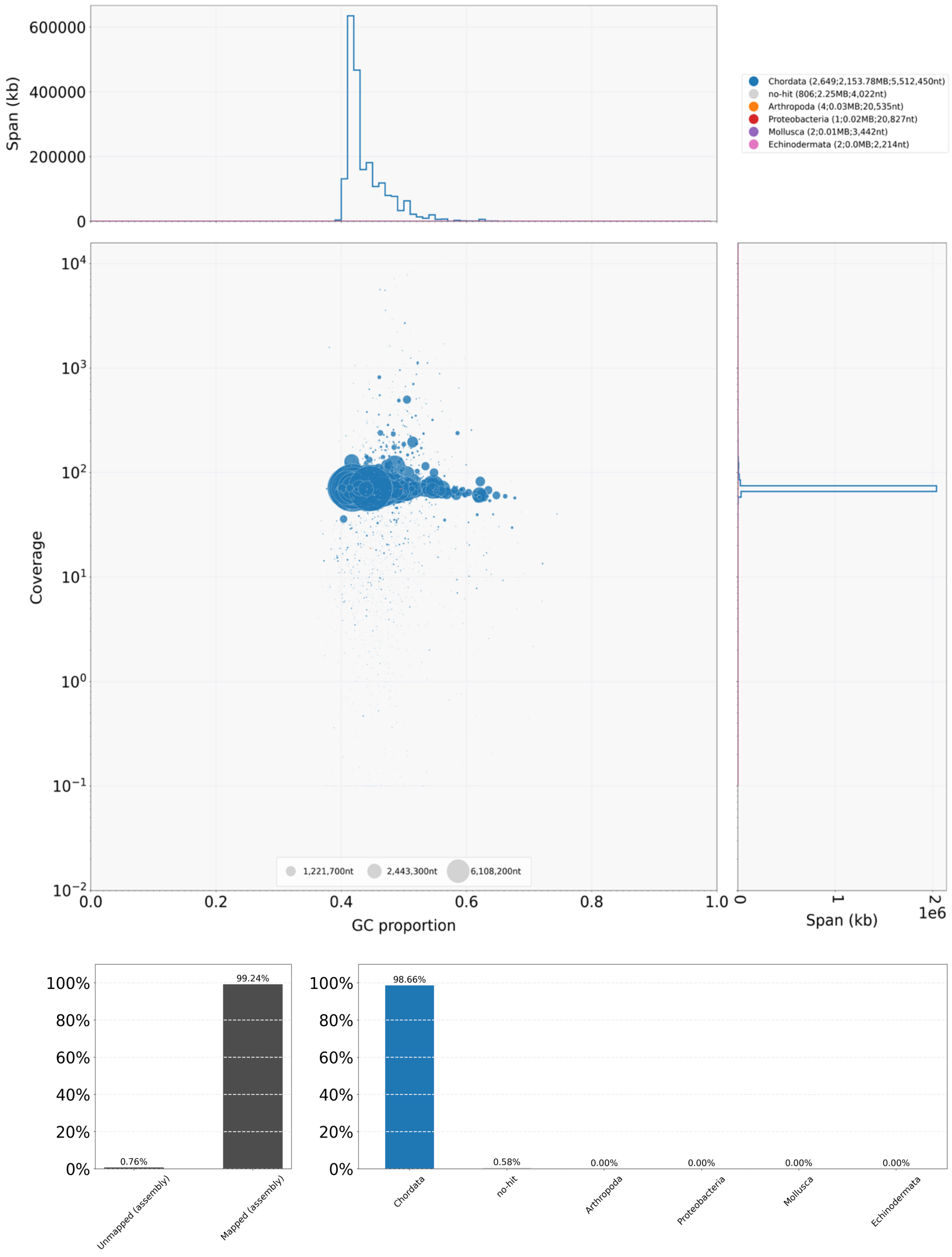


**Figure S3. GC-coverage blob-plot to evaluate assembly contamination.** Outputted by BlobTools v.1.1.1 (Laetsch and Blaxter, 2017) with our contig-level assembly. Scaffolds are coloured by phylum, and blob size represents scaffold length. Frequency histograms are plotted along the side of each axis. Lower bar charts show the proportion of scaffolds that were mapped to each phylum. Of the 99.24% of scaffolds that successfully mapped to our assembly, 98.66% of scaffolds mapped to Chordata (blue), and 0.58% produced no hit. This confirms minimal taxonomic contamination from other phyla in our assembly.

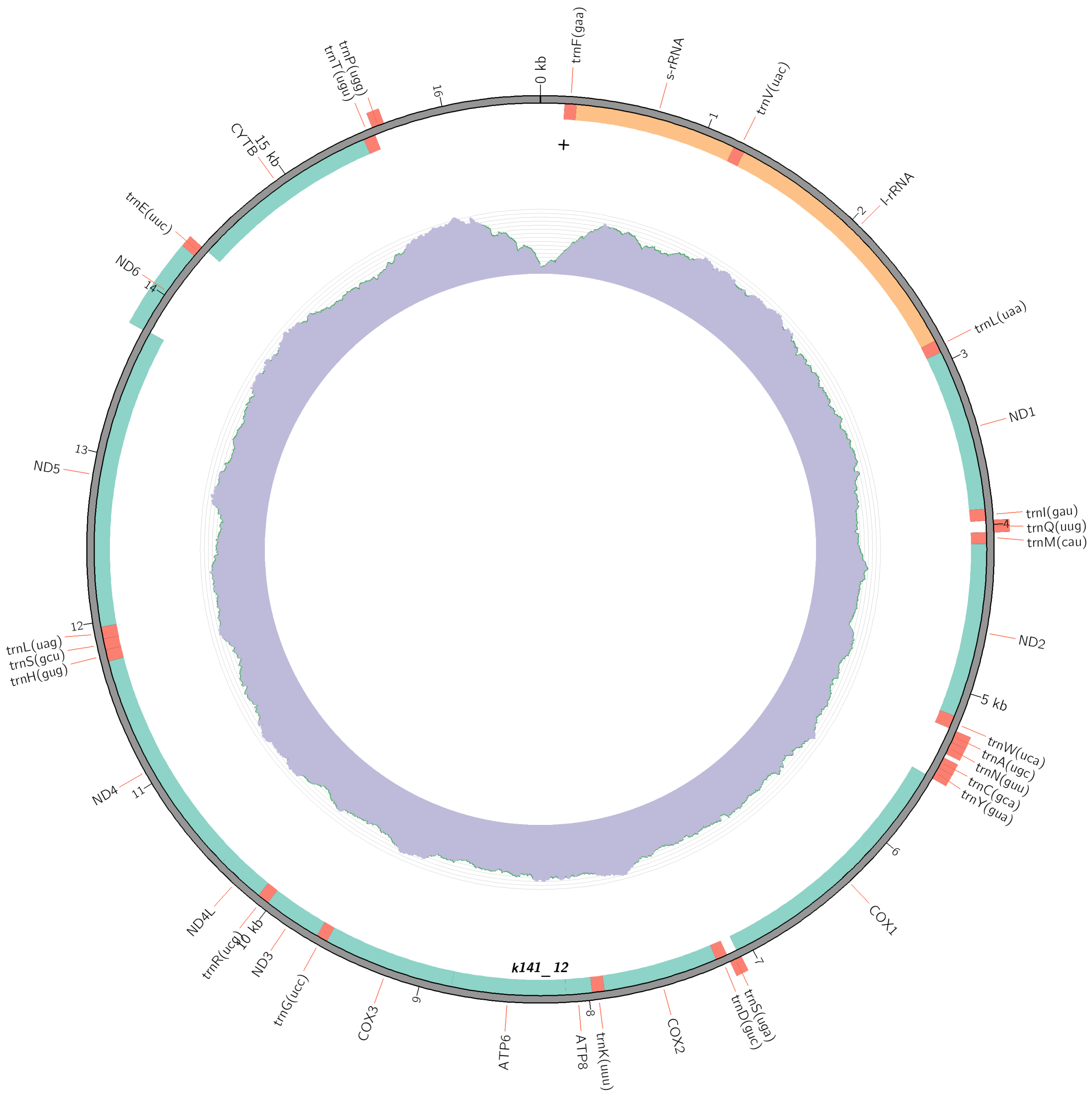


**Figure S4. Circos plot of our loggerhead mitochondrial assembly and annotation.** Length: 16,574 bp, GC content: 38.75%, total number of genes: 37 (protein coding genes: 13, tRNA genes: 22, rRNA genes: 2). The inner purple ring visualises coverage. Mitochondrial assembly, annotation and visualisation were performed with MitoZ v.3.4 (Meng *et al.*, 2019).

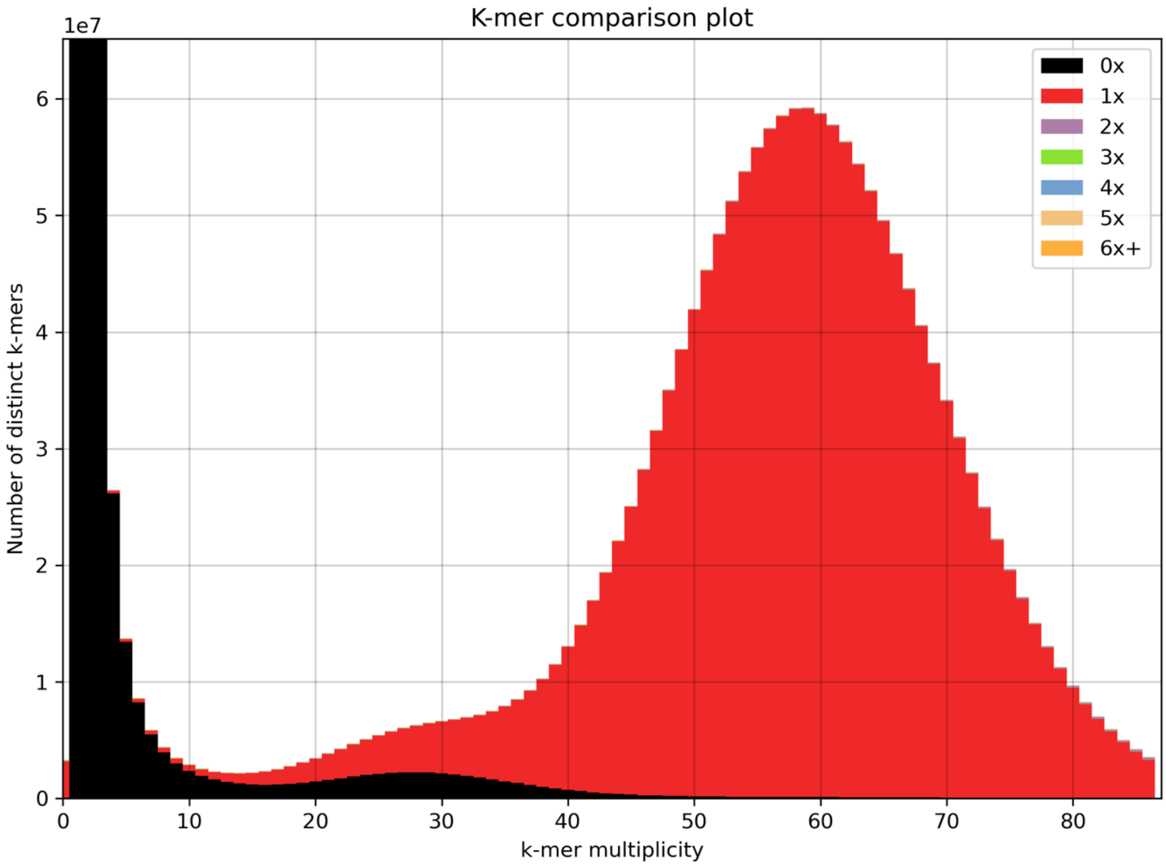


**Figure S5. K-mer spectrum plot of our loggerhead assembly.** Produced using KAT v.2.4.1 (Mapleson *et al.*, 2017), showing the frequency of k-mers in the assembly versus frequency of k-mers (i.e. sequencing coverage) in the raw Illumina reads. The first, black peak corresponds to k-mers present in the raw reads but missing from the assembly due to sequencing errors. The second, smaller peak corresponds to k-mers from heterozygous regions, and third peak corresponds to k-mers from homozygous regions. This plot confirms that our assembly is well haploidised.

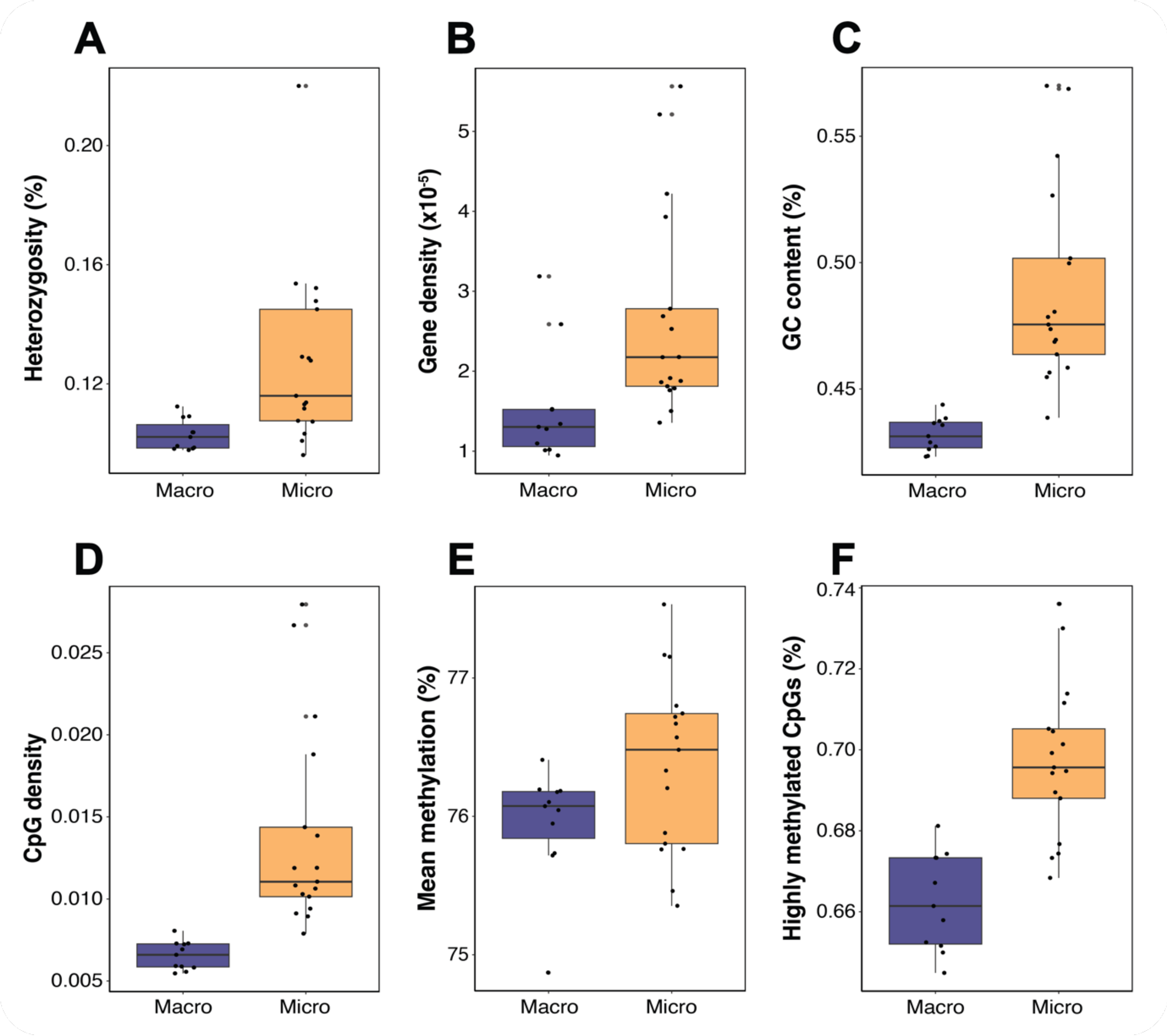


**Figure S6. Comparison of genome properties between macro- and microchromosomes.** Boxplots comparing genetic and epigenetic properties per chromosome, for the 11 macrochromosomes (dark blue) and 17 microchromosomes (yellow) of the loggerhead genome. Comparisons for: **(A)** heterozygosity (%), **(B)** gene density (total genes over chromosome length, x$10^{-5}$), **(C)** GC content (%), **(D)** CpG density (total CpGs over chromosome length), **(E)** methylation (%), and **(F)** proportion of highly methylated (>70%) CpGs (%). Microchromosomes are more heterozygous (W=29, p=0.002), gene-dense (W=28, p=0.001), GC-rich, CpG-dense (W=1, p<0.0001), and possess a larger proportion of highly methylated CpGs (W=54, p<0.0001).

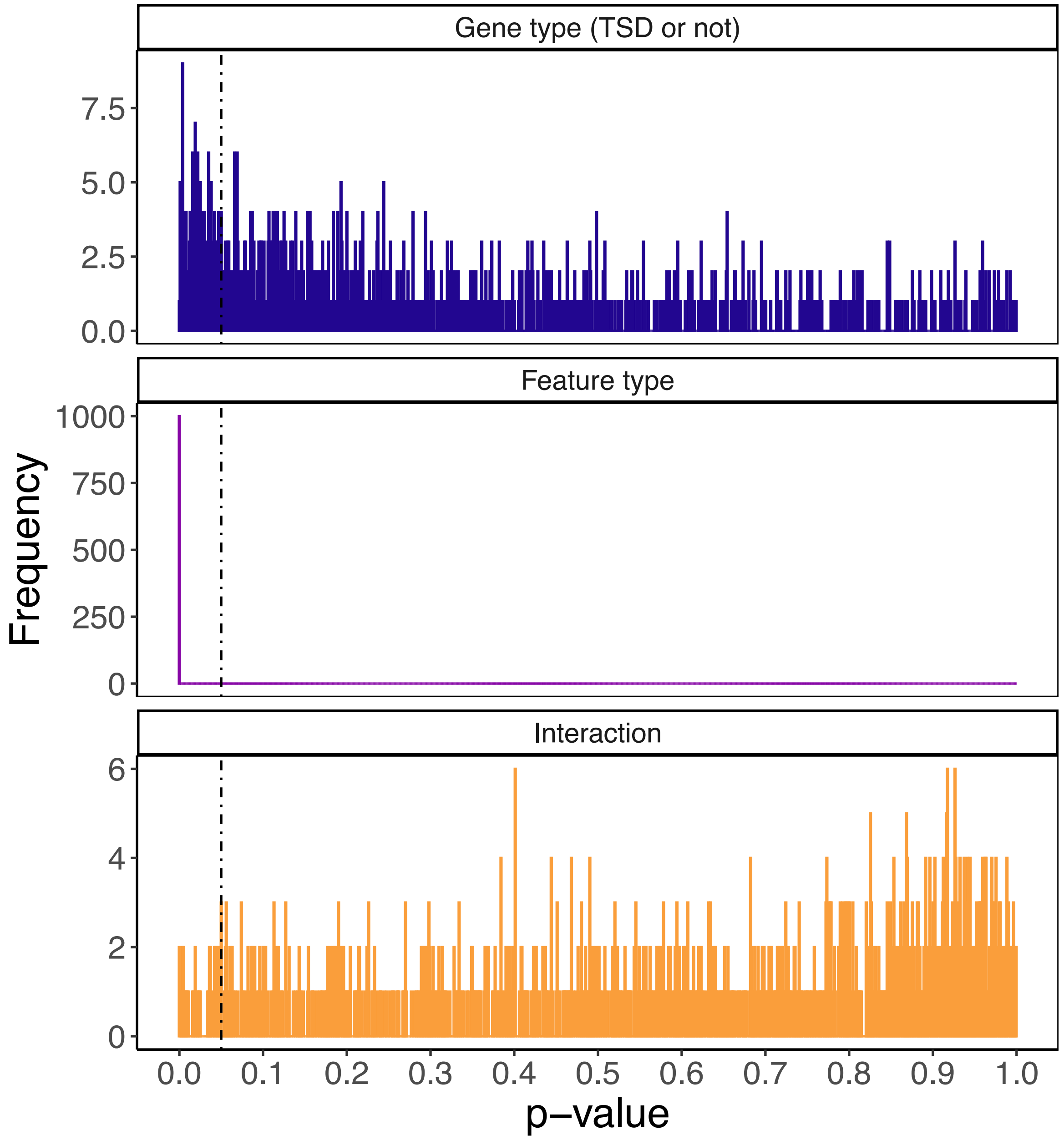


**Figure S7. P-value histogram from comparing mean methylation between TSD-linked genes and non-TSD-linked genes.** A linear model was used to test if mean methylation differed between 200 TSD-linked genes and 1000 random subsets of 200 non-TSD-linked genes, sampled from 15,041 single-copy orthologues shared between sea turtle species. Histograms are plotted for the p-values per iteration (n=1000), for each term in the ANOVA results table: gene category (TSD-linked versus non-TSD-linked, dark blue), genomic feature type (purple) and their interaction (Gene category x Feature type, yellow). The dashed line represents p=0.05. 155 (15.5%) tests passed p<0.05 for gene category, 1000 (100%) tests passed for feature type and 38 (0.38%) tests passed for the interaction term.

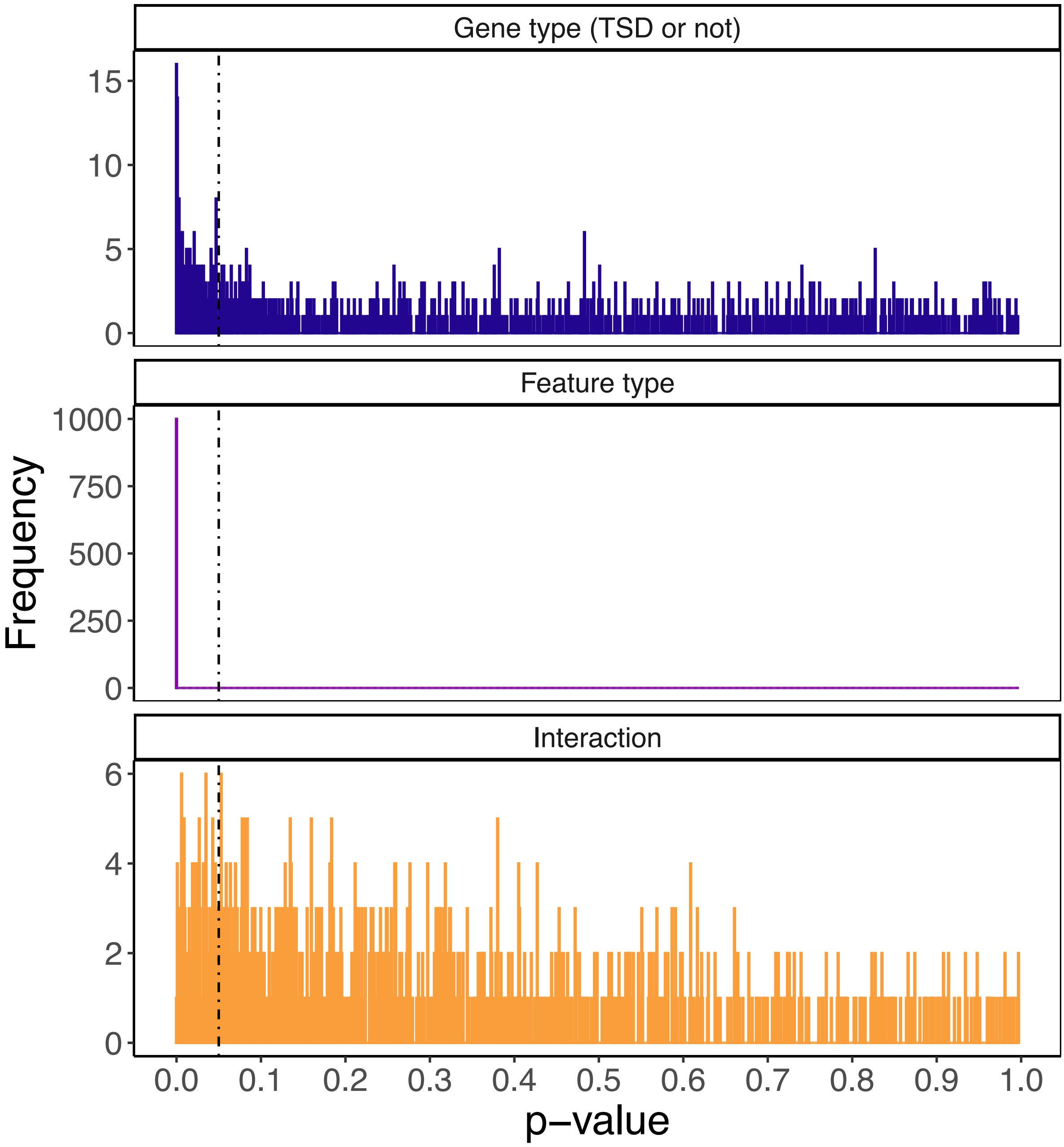


**Figure S7. P-value histogram from comparing the proportion of highly methylated CpGs between TSD-linked genes and non-TSD-linked genes.** A quasipoisson generalised linear model was used to test if the count of highly methylated CpGs differed between 200 TSD-linked genes and 1000 random subsets of 200 non-TSD-linked genes, sampled from 15,041 single-copy orthologues shared between sea turtle species. An offset of total CpG count was included in the model. Histograms are plotted for the p-values per iteration (n=1000), for each term in the ANOVA results table: gene category (TSD-linked versus non-TSD-linked, dark blue), genomic feature type (purple) and their interaction (Gene category x Feature type, yellow). The dashed line represents p=0.05. 189 (18.9%) tests passed p<0.05 for gene category, 1000 (100%) tests passed for feature type and 136 (1.36%) tests passed for the interaction term.
